# SIRT2 attenuates stress-induced skeletal muscle atrophy by inhibiting glucocorticoid receptor signalling

**DOI:** 10.1101/2025.10.22.683887

**Authors:** Ankit Kumar Tamta, Bhoomika Shivanaiah, Arathi Bangalore Prabhashankar, Sunayana Ningaraju, Seemadri Subhadarshini, Anurag Nagaraja Sharma, Harsha Mambully Jayaprakasan, Meera Krishnan E R, Apeksha Bhuyar, Devika Sunil, Amarjeet Shrama, Dimple Nagesh, Aastha Munjal, Sukanya Raghu, Manepalli Bhavya Madhuri, Dhevi Rajesh, Venketsubbu Ramasubbu, Mohsen Sarikhani, Himani Tandon, Perumal Arumugam Desingu, Latha Diwakar, Utpal Nath, Deepak Nair, Amit Singh, Ramanathan Sowdhamini, Narayanaswamy Srinivasan, Raul Mostoslavsky, Ninitha Asirvatham-Jeyaraj, Nagalingam Ravi Sundaresan

## Abstract

Skeletal muscle atrophy occurs in several diseases and is associated with chronic stress. Studies indicate that glucocorticoid receptor signalling is the major signalling pathway that mediates stress-induced muscle degeneration. Although the glucocorticoid signalling pathway is relatively well characterized, there is a need to identify modulators of this pathway that may be useful for drug targeting to ameliorate muscle atrophy. SIRT2 is a mammalian Sirtuin isoform known to mediate the longevity benefits of calorie restriction and exercise. Currently, the role of SIRT2 in regulating stress-induced skeletal muscle atrophy is unclear. Our study found that SIRT2 is a critical regulator of muscle homeostasis and is required to protect against stress-induced muscle atrophy. Interestingly, SIRT2 levels are reduced during glucocorticoid-induced muscle atrophy in mice. SIRT2 depletion exacerbates glucocorticoid-induced reduction in myotube diameter and atrophy gene expression. In contrast, SIRT2 overexpression ameliorates myotube atrophy in primary myotubes. Our findings indicate that SIRT2 knockout mice are susceptible to glucocorticoid-induced muscle atrophy, while muscle-specific SIRT2-transgenic mice exhibit improved muscle function and are protected from glucocorticoid-induced atrophy. Mechanistically, SIRT2 binds to the glucocorticoid receptor to negatively regulate its activity, possibly via deacetylation of critical residues in its DNA-binding domain. Our findings suggest that SIRT2 activation may protect against glucocorticoid-induced skeletal muscle atrophy and serve as a potential therapeutic target for treating muscle atrophy.

## INTRODUCTION

Skeletal muscles constitute nearly 40% of the body weight and are among the largest tissues in the human body. They are our body’s largest protein reservoirs, storing 50-70% of the total protein [1]. Loss of muscle homeostasis results in muscle atrophy, a debilitating disorder that increases mortality and reduces the quality of life [2, 3]. Muscle atrophy is observed to develop during various ageing-associated diseases, including cardiovascular diseases (CVDs), renal failure, arthritis, cancer, and diabetes [4–9]. Furthermore, several forms of chronic stress, including starvation and immobilization, are also known to induce muscle atrophy [10].

Muscle atrophy is characterized by a reduction in myofiber size, resulting in the loss of muscle mass [11]. Under normal physiological conditions, muscle mass and function are maintained by regulating protein anabolism and catabolism. This allows the muscle to adapt to changes in nutrient availability and physical activity [12]. A critical molecular pathway involved in muscle protein homeostasis is the Akt-mTOR pathway, which is regulated in response to a variety of upstream signaling molecules, including glucocorticoids, growth hormone (GH), insulin-like growth factor-1 (IGF1), insulin, epinephrine, and testosterone [13–16]. Specifically, stress-induced skeletal muscle atrophy is primarily mediated by glucocorticoid signaling. As an adaptive response to stress, cortisol is released into the bloodstream and transported to target organs and tissues, including skeletal muscles, where it binds to and activates the cytosolic glucocorticoid receptor. In its active form, the Glucocorticoid Receptor (GR) translocates into the nucleus, binds to the DNA, and regulates the target gene expression [17]. Glucocorticoid signaling inhibits protein synthesis and promotes protein breakdown by various mechanisms, ultimately resulting in atrophy [18]. It inhibits the transport of amino acids into muscles, resulting in halting of protein synthesis [19]. It also suppresses insulin and IGF-1 signaling to inhibit protein synthesis. Notably, translational repressor 4E-BP1 and ribosomal S6 kinase 1 (S6K1) are dephosphorylated in response to glucocorticoids, inhibiting protein translation [20–23]. Glucocorticoid signaling also promotes proteolysis by activating proteolytic systems, including autophagy, the ubiquitin-proteasome system (UPS), and calcium-dependent calpains [24]. Glucocorticoids activate the expression of UPS components and trigger ubiquitination of the targeted proteins [25]. Additionally, glucocorticoids can also directly degrade proteins through different 20S proteasome subunits [26]. Together via these mechanisms, chronic stress-induced glucocorticoid signaling inhibits protein synthesis and enhances protein degradation, eventually leading to muscle degradation and atrophy - severely compromising the lifespan and health span of individuals. Therefore, identifying modulators of the GR signaling-induced skeletal muscle atrophy that can serve as therapeutic targets for treating muscular atrophy is essential.

Calorie-restriction-based dietary and exercise-based activities have been shown to prevent the development of muscle atrophy. It is well established that the health benefits of calorie restriction are primarily mediated by the NAD^+^-dependent class III histone deacetylases, Sirtuins (SIRTs). Mammals express seven isoforms of NAD^+^-dependent sirtuins (SIRT1-7), each of which localizes to different subcellular compartments and regulates critical biological processes, including longevity, metabolism, gene expression, DNA repair, and inflammation [27–29]. Importantly, SIRT2, the mammalian ortholog of the yeast-origin gene Hst2, has been demonstrated to mediate the longevity benefits of calorie restriction [30]. Accordingly, SIRT2 expression is reported to be upregulated during calorie restriction in mice [31]. SIRT2 also regulates critical transcription factors, including forkhead transcription factors of class O, namely FOXO1 and FOXO3, in response to nutrient deprivation to promote lipolysis and maintain energy homeostasis in mice [31, 32]. Our previous study indicates that SIRT2 inhibits the activity of transcription factor NFATc2 to promote cardiac homeostasis in mice [33]. SIRT2 is expressed abundantly in skeletal muscles, where it has been reported to alleviate stress-induced atrophy *in vitro* in C2C12 myotubes by inhibiting autophagy [34]. Other studies have also demonstrated a role for SIRT2 as a negative regulator of insulin signalling *in vitro* [35]. These studies indicate a potential role for SIRT2 in regulating muscle structure and function. However, due to a lack of appropriate animal models for stress-induced skeletal muscle atrophy research, the role of SIRT2 in regulating skeletal muscle dynamics *in vivo* remains elusive. Furthermore, the molecular mechanisms by which SIRT2 may regulate skeletal muscle homeostasis are poorly understood.

In the current study, we address these gaps by investigating the role of SIRT2 in the regulation of stress-induced skeletal muscle atrophy using established *in vitro* models such as murine primary myotubes and animal models like SIRT2 knockout mice, as well as a muscle-specific SIRT2-transgenic mouse. Interestingly, our results indicate that the expression of SIRT2 is downregulated in skeletal muscles under stress conditions induced upon treatment with the GR agonist dexamethasone. Importantly, we found that SIRT2-overexpressing mice are protected from GR-induced muscle atrophy. Our results suggest that SIRT2 ameliorates dexamethasone-induced muscle atrophy, potentially by interacting with and deacetylating novel residues in GR to inhibit its DNA binding activity. Overall, our study implicates a protective role for SIRT2 against stress-induced muscle atrophy.

## RESULTS

### SIRT2 expression is downregulated in *in vitro* and *in vivo* models of glucocorticoid-induced muscle degeneration

Dexamethasone (dex) is a synthetic glucocorticoid analog that is administered as an anti-inflammatory drug for the treatment of a variety of diseases [36]. However, its systemic use is linked with the development of muscle atrophy [37]. Based on this finding, dex has been adapted to induce skeletal muscle atrophy in *in vitro* and *in vivo* model systems [38]. Thus, to generate an animal model for investigating stress-induced muscle atrophy *in vivo*, we injected four-month-old C57BL/6J mice with dex for 15 days at a 24-hour interval. Post-dex treatment, we observed a significant reduction in mouse body weight (**Figure 1A, B**). Furthermore, dex-treated mice were also characterized by reduced muscle weight to tibia length ratio in type II fiber dominant gastrocnemius and tibialis anterior muscles (**Figure 1C**).

**Figure 1.**
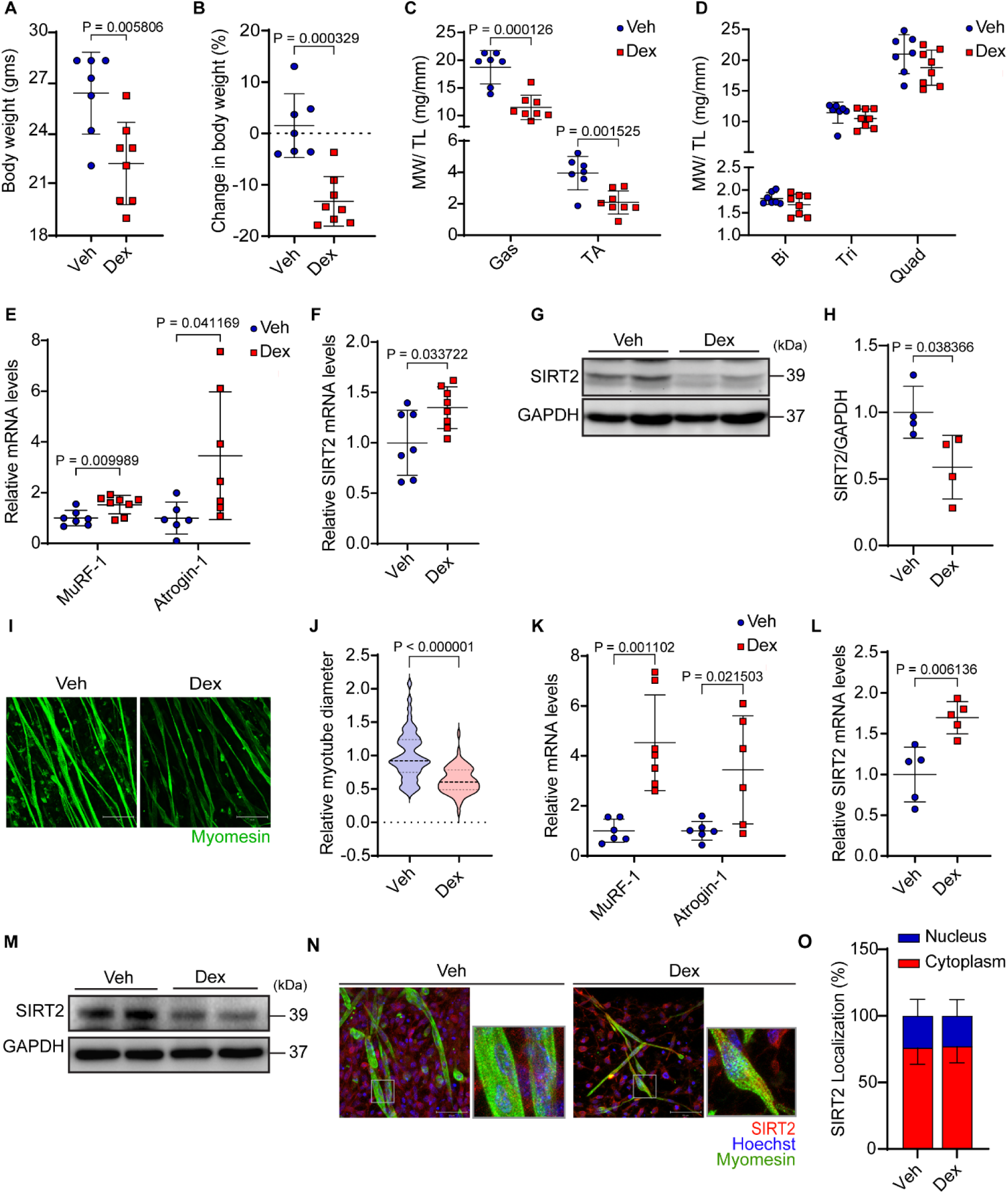
SIRT2 levels in stress-mediated muscle atrophy. **(A)** The Body weight measurements of wild-type mice after 15 days of dex injection (10mg/kg/day) compared to the vehicle group. Data represented as mean ± s.d., *n* = 7-8. **(B)** Change in percentage body weight of mice post 15 days of dex injection (10mg/kg/day) compared to vehicle group. Data represented as mean ± s.d., *n* = 7-8. **(C)** Muscle weight to tibia length ratio of the gastrocnemius and tibialis anterior muscles in mice administered with dex or vehicle. Data represented as mean ± s.d., *n* = 7-8. **(D)** Muscle weight to tibia length ratio of the biceps (Bi), triceps (Tri), and quadriceps (Quad) muscles in mice administered with dex or vehicle. Data represented as mean ± s.d., *n* = 7-8. **(E)** qRT-PCR analysis for the relative expression of MuRF-1 and Atrogin-1 in the gastrocnemius muscle of mice injected with either vehicle or dex. Data represented as mean ± s.d., *n* = 6-8. **(F)** qRT-PCR analysis for the relative expression of SIRT2 in the gastrocnemius muscle of mice injected with either vehicle or dex. Data represented as mean ± s.d., *n* = 7-8. **(G)** Representative immunoblot of the gastrocnemius muscles of mice injected with dex. Indicated sizes are in kilodaltons (kDa). **(H)** Quantification of SIRT2 levels in the gastrocnemius muscles of dex-treated mice of Figure 1(G). Protein levels were normalized with GAPDH. Data represented as mean ± s.d., *n* = 4. **(I)** Representative immunofluorescence images of the primary rat myotubes treated either dex (50µM) or vehicle, stained with antibodies against myomesin (green). Scale bar = 50 µm. **(J)** Quantification of relative myotube diameters of the dex-treated primary rat myotubes normalized with the vehicle. Data represented as median with 25 and 75 percentile *n* = 3, N= 56-64. **(K)** qRT-PCR analysis for the relative expression of MuRF-1 and Atrogin-1 in the primary rat myotubes treated with dex and normalized with the vehicle. Data represented as mean ± s.d., *n* = 6-7. **(L)** qRT-PCR analysis for the relative expression of SIRT2 in the vehicle and dex-treated primary rat myotubes. Data represented as mean ± s.d., *n* = 5. **(M)** Representative immunoblot showing SIRT2 levels in the dex-treated primary rat myotubes. Indicated sizes are in kilodaltons (kDa). **(N)** Representative immunofluorescence images showing SIRT2 localization in the vehicle and dex-treated primary rat myotubes. Myotubes were stained with antibodies against myomesin (green), Hoechst (blue), and SIRT2 (red). SIRT2 intensity has been modified to represent its localization better. Scale bar = 50 µm. **(O)** Quantification of SIRT2 localization upon vehicle and dex-treated conditions in primary rat myotubes. *n* = 3. The two-tailed unpaired student’s t-test with Welch’s correction was used for statistical analysis (A, B, F, H, J, L). Unpaired t-tests per row with Holm-Sidak’s multiple comparisons test was used for (C, D) Two-way ANOVA with Sidak’s multiple comparisons test (E, K) Kruskal-Wallis test with Dunn’s multiple comparisons (O) was used for statistical analysis.

Consistent with previous reports indicating Type II muscle fiber-specific action of dex, we did not observe any significant difference in weights of type I fiber dominant muscles (**Figure 1D**) [39, 40]. Similarly, we observed an upregulated mRNA expression of atrophy markers, such as MuRF-1 and Atrogin-1, in the gastrocnemius of the dex-injected mice (**Figure 1E**). These results indicate that dex injection induced muscle atrophy *in vivo* in our model system. A recent study demonstrated that SIRT2 expression is reduced upon dexamethasone treatment in mice skeletal muscle [34]. To investigate the role of SIRT2 in stress-induced muscle atrophy, we evaluated SIRT2 expression in the skeletal muscles of dex-injected mice. Interestingly, we found that, compared to control mice, the SIRT2 mRNA levels were significantly upregulated in the gastrocnemius of dex-treated mice (**Figure 1F**). However, the SIRT2 protein expression was significantly downregulated in these mice (**Figure 1G, H**).

We treated primary myotubes with dex to further validate these results *in vitro*. Similar to the results obtained in the *in vivo* model, we observed a substantial decrease in the diameter of the myotubes upon dex treatment (**Figure 1I, J**). Additionally, dex-treated myotubes showed an increase in the expression of the atrogenes, MuRF-1, and Atrogin-1, indicating atrophy (**Figure 1K**). Similar to our *in vivo* results, these dex-treated myotubes also displayed increased SIRT2 mRNA expression (**Figure 1L**) and reduced protein expression compared to the untreated control myotubes (**Figure 1M**). To account for the reduced SIRT2 protein levels upon dex treatment, we used MG132, a proteasome inhibitor, and observed that dex-associated reduction of SIRT2 was mitigated upon proteasome inhibition **(Figure S1A-C)**. This suggests that dex-associated SIRT2 depletion may be due to increased ubiquitination of SIRT2 and subsequent proteasomal degradation, and that enhanced mRNA expression of SIRT2 may be a compensatory response for reduced SIRT2 protein levels. SIRT2 has been reported to shuttle between the cytoplasm and the nucleus in response to various stress stimuli [41, 42]. Therefore, we evaluated the effect of dex treatment on the subcellular localization of SIRT2 and found that SIRT2 remains localized primarily in the cytoplasm of myotubes (**Figure 1N, O**), indicating that dex treatment did not affect the subcellular localization of SIRT2. Together, these results suggest a potential role for SIRT2 in developing dexamethasone-induced muscle atrophy.

### SIRT2 regulates glucocorticoid-induced myotube atrophy *in vitro*

Since our findings suggest a potential role for SIRT2 in stress-induced muscle atrophy, we developed *in vitro* model systems to investigate the role of SIRT2 in dex-induced myotube atrophy. First, we generated a SIRT2 knockdown myotube population using a siRNA-based approach to study the effect of SIRT2 depletion in primary myotubes (**Figure 2A, B**). SIRT2-depleted myotubes were characterized by reduced myotube diameter compared to the control myotubes, and their myotube diameter was further reduced upon dex treatment (**Figure 2C, D**). Moreover, dex-treated SIRT2 knockdown myotubes also showed enhanced mRNA expression of atrogin-1 and MuRF-1 compared to the wild-type dex-treated myotubes. However, SIRT2 depletion alone did not induce atrogene expression in these myotubes under basal conditions (**Figure 2E**). These results indicate that SIRT2 depletion exacerbates stress-induced muscle atrophy in a cell-autonomous manner. To validate these results further, we overexpressed SIRT2 in primary myotubes (**Figure 2F, G**) and observed that SIRT2 overexpressing myotubes were also resistant to dex-induced reduction in myotube diameter (**Figure 2H, I**). Similarly, compared to the wild-type control, the SIRT2 overexpressing myotubes also showed a marked reduction in the expression of MuRF-1 and atrogin-1 under dex-treated conditions (**Figure 2J, K**). These findings suggest that SIRT2 may play a protective role in stress-induced muscle atrophy.

**Figure 2.**
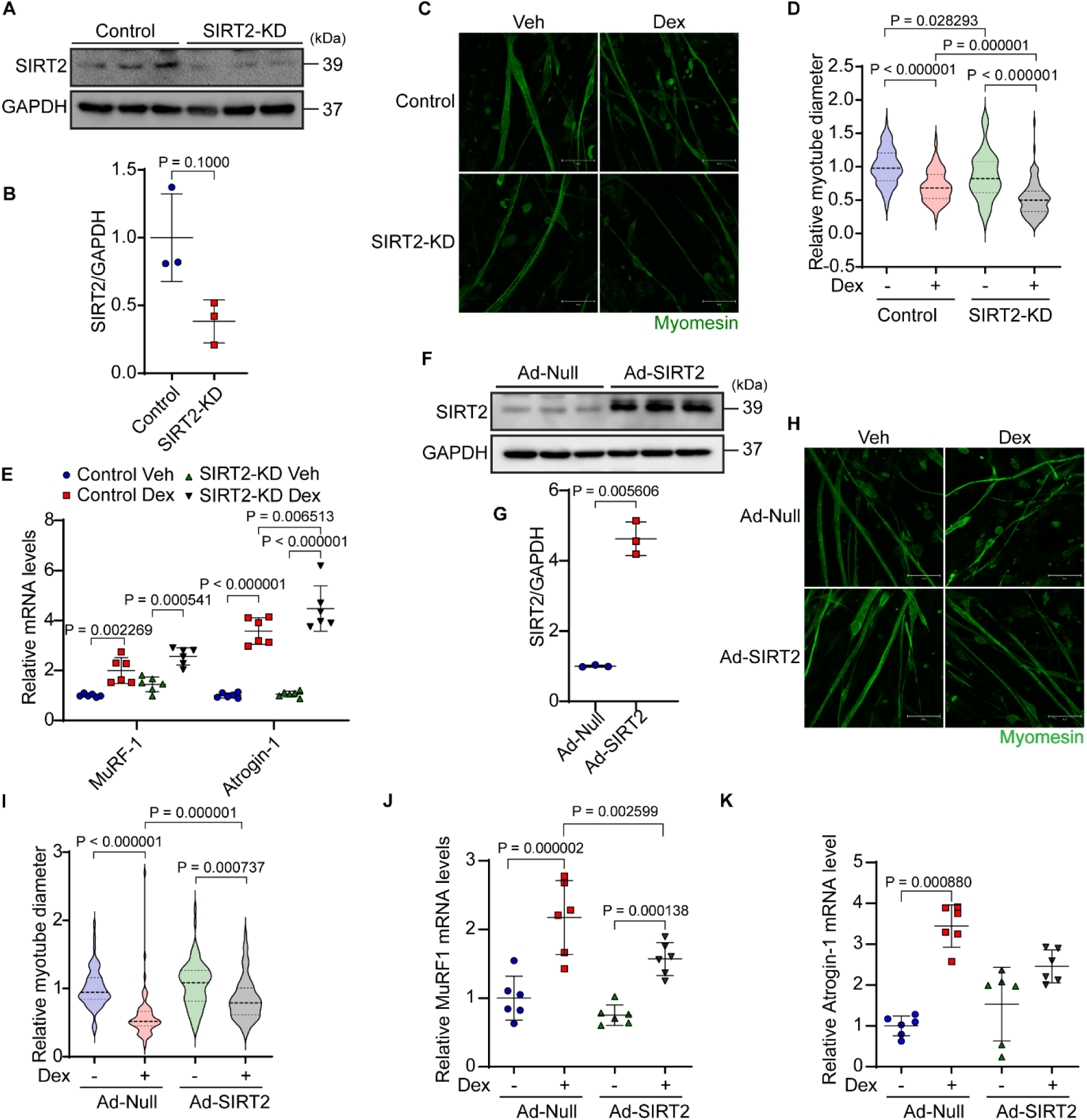
SIRT2 regulates myofiber atrophy in an *in vitro* atrophy model. **(A)** Representative immunoblot to verify SIRT2 depletion in primary myotubes. Indicated sizes are in kilodaltons (kDa). **(B)** Quantification of SIRT2 levels using immunoblot shown in Figure 2(A). Protein levels were normalized with GAPDH. Data represented as mean ± s.d., *n* = 3. **(C)** Representative immunofluorescence images showing diameter of the myotubes in control and SIRT2 depleted conditions when treated with vehicle or dex. Myotubes were stained with antibodies against myomesin (green). Scale bar = 50 µm. **(D)** Quantification of relative myotube diameters in dex-treated primary myotubes under SIRT2 depleted conditions, normalized with the vehicle-treated control. Data represented as median with 25 and 75 percentile *n* = 3, N= 58-122. **(E)** qRT-PCR analysis for the relative expression of MuRF-1 and Atrogin-1 in SIRT2 depleted myotubes under dex treatment, normalized with vehicle-treated controls. Data represented as mean ± s.d., *n* = 6. **(F)** Representative immunoblot showing adenovirus-mediated SIRT2 overexpression in the primary myotubes. Indicated sizes are in kilodaltons (kDa). **(G)** Quantification of SIRT2 levels using immunoblot shown in Figure 2(F). Protein levels were normalized with GAPDH. Data represented as mean ± s.d., *n* = 3. **(H)** Representative immunofluorescence images showing diameter of the myotubes in null and SIRT2 overexpression conditions when treated with either vehicle or dex. Myotubes were stained with antibodies against myomesin (green). Scale bar = 50 µm. **(I)** Quantification of relative myotube diameters in dex-treated primary myotubes under adenovirus-mediated SIRT2 overexpression condition, normalized with the vehicle-treated null adenovirus condition. Data represented as median with 25 and 75 percentile *n* = 3, N= 68-75. **(J)** qRT-PCR analysis for the relative expression of MuRF-1 in the SIRT2 overexpressing myotubes under dex treatment, normalized with vehicle-treated null adenovirus condition. Data represented as mean ± s.d., *n* = 6. **(K)** qRT-PCR analysis for the relative expression of Atrogin-1 in the SIRT2 overexpressing myotubes under dex treatment, normalized with vehicle-treated null adenovirus condition. Data represented as mean ± s.d., *n* = 6. Two-tailed Mann-Whitney statistical test (B) Kruskal-Wallis test with Dunn’s multiple comparisons (D, I, K) Two-way ANOVA with Tukey’s multiple comparisons test was used for statistical analysis (E, J). The two-tailed unpaired student’s t-test with Welch’s correction (G) was used for statistical analysis.

### SIRT2 knockout mice are susceptible to glucocorticoid-induced skeletal muscle atrophy

Since our previous results suggest a protective role for SIRT2 in the development of muscle atrophy, we developed animal models to characterize the role of SIRT2 in the development of skeletal muscle atrophy *in vivo*. To further validate our previous results indicating that SIRT2 expression is downregulated in the skeletal muscles of dex-treated mice, we utilized a whole-body SIRT2 knockout (SIRT2^-/-^) mouse model (**Figure 3A**). We assessed the glucose and insulin sensitivity of SIRT2^-/-^ mice. We observed that SIRT2 knockout mice have a better glucose clearance rate and are more sensitive to insulin response than wild-type mice **(Figure S2A-H)**. We then isolated and cultured myotubes from these SIRT2^-/-^ mice and treated them with dex. Consistent with our previous models, we observed a significant reduction in myotube diameter upon dex treatment, which was further exacerbated in SIRT2^-/-^ mice (**Figure S2A-C)**. Next, to understand the role of SIRT2 in stress-induced changes in muscle physiology *in vivo*, we treated control and SIRT2 knockout mice with dex and observed changes in physiological parameters. We observed that SIRT2^-/-^ mice exhibit a significant reduction in body weight compared to wild-type control mice upon dex treatment (**Figure 3B, S2D)**. Further, the reduction in gastrocnemius and tibialis anterior muscle weight was also pronounced in SIRT2^-/-^ mice compared to the wild-type control mice (**Figure 3C, D, S2E, F)**. Meanwhile, the weight of other muscles largely remained unchanged, validating the type II muscle-specific action of glucocorticoids on muscles **(Figure S2G-L)**. Interestingly, the cross-sectional area of gastrocnemius muscle fibers was significantly reduced in SIRT2^-/-^ mice compared to the wild-type mice treated with dex (**Figure 3E, F**). These mice also displayed an overall increase in the number of small cross-sectional area myofibres in the gastrocnemius muscles (**Figure 3G**). Moreover, muscles from these mice exhibited increased fibrosis compared to the wild-type controls (**Figure 3E, H**). Consistent with these observations, we also found upregulated atrogenes expression in dex-treated SIRT2^-/-^ mice compared to the wild-type mice (**Figure 3I**).

**Figure 3.**
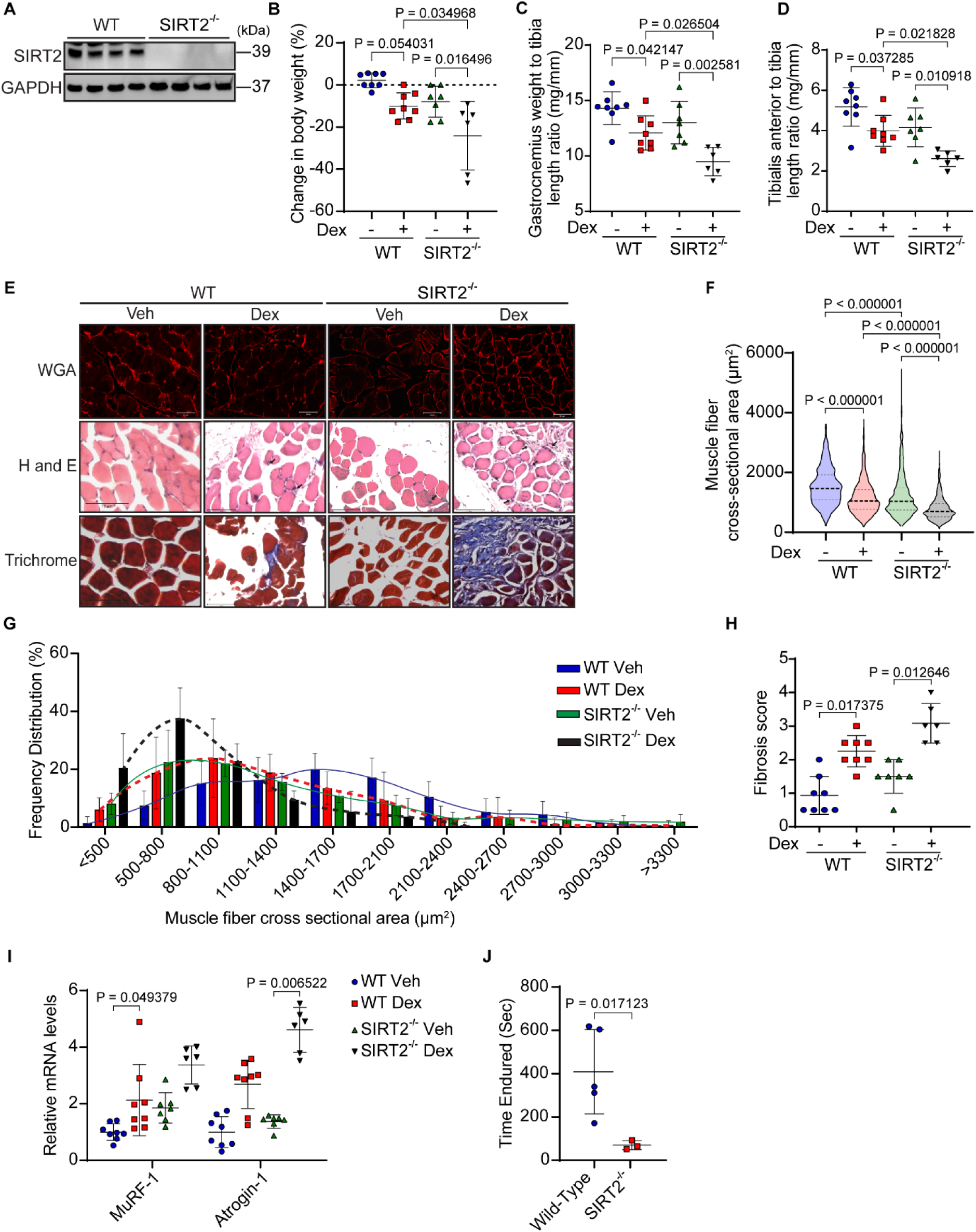
SIRT2^-/-^ mice develop skeletal muscle atrophy. **(A)** Representative immunoblot to verify SIRT2 deficiency in SIRT2^-/-^ mice. Indicated sizes are in kilodaltons (kDa). **(B)** Percentage change in mice’s body weight post 15 days of injection with dex(10mg/kg/day) compared to the vehicle group in wild-type and SIRT2^-/-^ mice. Data represented as mean ± s.d., n = 6-8. **(C)** Muscle weight to tibia length ratio of the gastrocnemius in wild-type and SIRT2^-/-^ mice injected with either vehicle or dex(10mg/kg/day). Data represented as mean ± s.d., n = 6-8. **(D)** Muscle weight to tibia length ratio of the tibialis anterior in wild-type and SIRT2^-/-^ mice injected with either vehicle or dex(10mg/kg/day). Data represented as mean ± s.d., n = 6-8. **(E)** Representative cross sections showing the gastrocnemius muscles of wild-type and SIRT2^-/-^ mice injected with either vehicle or dex. Sections were stained using WGA, H and E, and Trichrome staining to depict muscle fiber cross sectional area, morphology, and fibrosis, respectively. Scale bars = 50, 75, and 75 µm, respectively. **(F)** Quantification of WGA images represented in Figure 3(E), showing muscle fiber cross sectional area of gastrocnemius muscles in wild type and SIRT2-/- mice administered with either vehicle or dex. Data are shown as median with 25 and 75 percentiles. n = 3-4, N = 548-1019. **(G)** Quantification of WGA images represented in Figure 3(E), showing the frequency distribution of cross sectional fiber area of gastrocnemius muscles in wild type and SIRT2^-/-^ mice administered with either vehicle or dex. Data are shown as mean ± s.d., n = 3-4. **(H)** Quantification of Masson’s Trichrome images represented in Figure 3(E), showing fibrosis score of gastrocnemius muscles in wild type and SIRT2-/- mice administered with either vehicle or dex. Scoring was done in a blind fashion. Data are shown as mean ± s.d., n = 6-8. **(I)** Relative expression of atrogenes (MuRF-1 and Atrogin-1) in the gastrocnemius muscles of SIRT2-/- mice under the vehicle or dex treatment, normalized with the vehicle-treated wild-type control. Data are shown as mean ± s.d., n = 6-8. **(J)** Quantification of time the wild-type and SIRT2^-/-^ mice endured in the rotating rod. Data are shown as mean ± s.d., n = 3-5. Two-way ANOVA with Tukey’s multiple comparisons (B, C, D) Kruskal-Wallis test with Dunn’s multiple comparisons (F, H, I) The two-tailed unpaired student’s t-test with Welch’s correction (J) was used for statistical analysis.

To understand whether these changes in muscle physiology and molecular signalling affect muscle strength and function in SIRT2^-/-^ mice, we performed a rotarod performance test using control and SIRT2^-/-^ mice [43]. We trained the mice to run on the rotarod instrument and found that at the basal level, SIRT2^-/-^ mice had reduced muscle performance compared to the control (**Figure 3J, S2M)**. We then assessed the muscle fiber type composition in the gastrocnemius muscles of SIRT2^-/-^ mice when treated with dex. In SIRT2^-/-^ mice, we observed a greater content of Type I and Type IIa muscles at the basal level than wild-type mice. However, their content in SIRT2^-/-^ mice was reduced upon dex treatment **(Figure S3A-F)**. These results suggest that SIRT2 deficiency exacerbates glucocorticoid-induced muscle atrophy, potentially influencing the fiber distribution pattern *in vivo*.

### Muscle-specific SIRT2-transgenic mice are protected from glucocorticoid-induced skeletal muscle atrophy

Since our previous findings indicate that overexpression of SIRT2 *in vitro* protects myotubes from dex-induced atrophy, we hypothesized that SIRT2-overexpressing mice might be protected from dex-induced skeletal muscle atrophy. To evaluate this, we generated novel muscle-specific SIRT2-transgenic mice (msSIRT2-Tg) (**Figure 4A, B**). First, we characterized these mice for various physiological parameters to assess the potential effects of SIRT2 overexpression on mice, if any. These mice appeared healthy and devoid of visible defects. Further, they displayed no significant difference in body weight compared to the non-transgenic control mice (N-Tg) (**Figure 4C**). We then assessed various blood parameters in these mice, including RBC size and count, hemoglobin concentration, leukocyte, neutrophils, lymphocyte, monocyte, and platelets count. We found no significant differences in these parameters between msSIRT2-Tg mice and the N-Tg control mice **(Figure S4A-K, S5A-K).** The msSIRT2-Tg mice also had normal liver and kidney function profiles **(Figure S4L-Y, S5L-X)** and plasma ion homeostasis **(Figure S4Z-AB)**. These mice also displayed normal glucose and insulin clearance rates **(Figure S6A-H),** suggesting that msSIRT2-transgenic mice display no overt symptoms of diabetes. Together, these findings suggest that increased SIRT2 expression in skeletal muscles does not affect the normal physiological conditions in mice.

**Figure 4.**
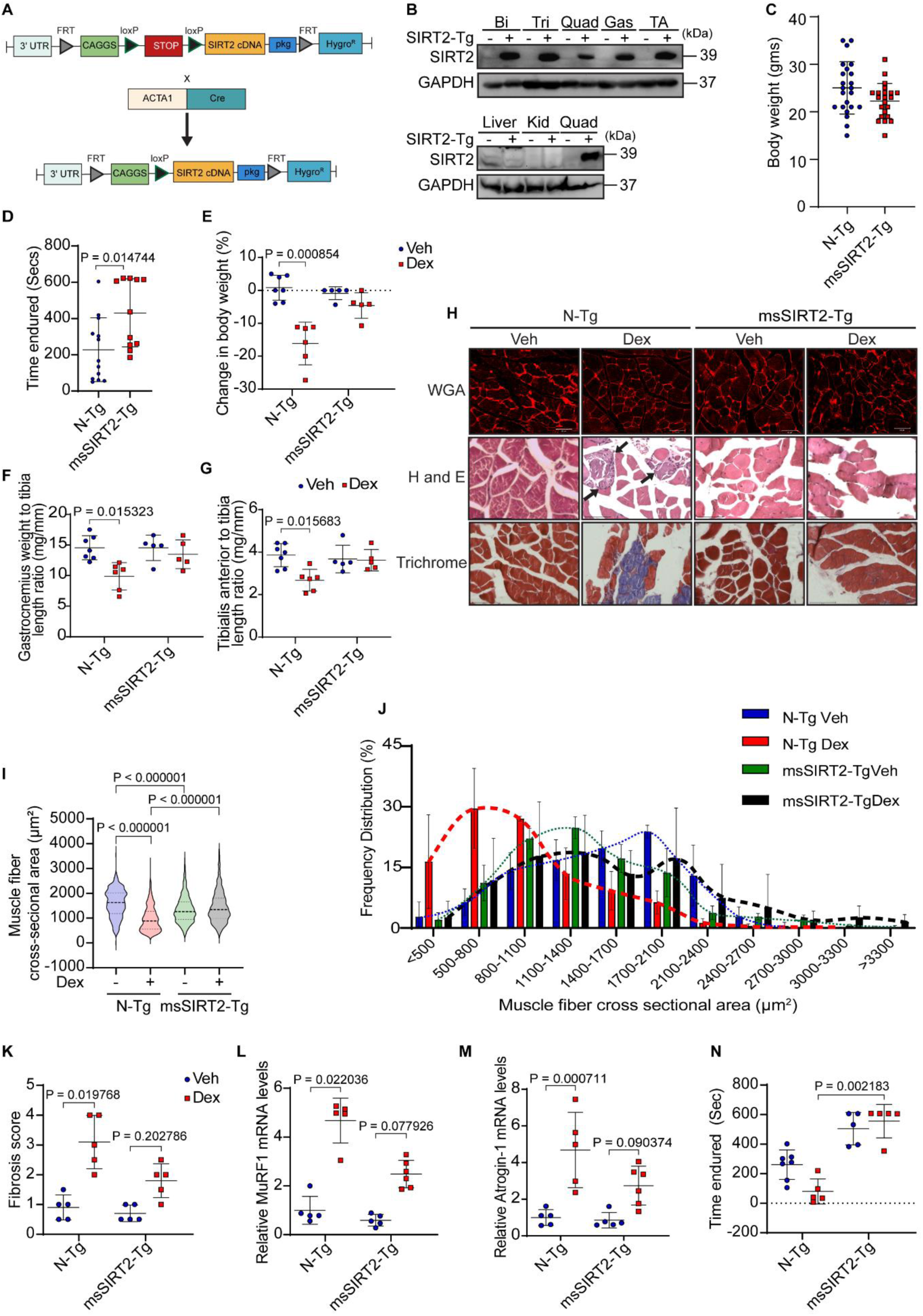
msSIRT2-transgenic mice are protected from skeletal muscle atrophy. **(A)** Schematic diagram representing the generation of the muscle-specific SIRT2-transgenic mice line. **(B)** Representative western blot images showing SIRT2 overexpression in various muscles (top) biceps (Bi), triceps (Tri), quadriceps (Quad), gastrocnemius (Gas), and tibialis anterior (TA) and different organs (below) liver, kidney (kid)and quadriceps (Quad) in msSIRT2-transgenic and non-transgenic mice. Indicated band sizes are in kilodaltons (kDa). **(C)** Graphical representation of body weight of msSIRT2 transgenic (male) compared to non-transgenic mice (males). Data represented as mean ± s.d., n = 22-23. **(D)** Quantification of time the msSIRT2-transgenic and non-transgenic mice endured in the rotating rod. Data are shown as mean ± s.d. n = 11-13. **(E)** Percentage change in mice’s body weight post 15 days of injection with dex(10mg/kg/day) compared to the vehicle group in non-transgenic and msSIRT2-transgenic mice. Data represented as mean ± s.d. n = 5-7. **(F)** Gastrocnemius weight to tibia length ratio in non-transgenic and msSIRT2-transgenic mice injected with either vehicle or dex(10mg/kg/day). Data represented as mean ± s.d. n = 5-7. **(G)** Tibialis anterior weight to tibia length ratio in non-transgenic and msSIRT2-transgenic mice injected with either vehicle or dex(10mg/kg/day). Data represented as mean ± s.d. n = 5-7. **(H)** Representative images showing the cross-sectional sections of the gastrocnemius muscles of non-transgenic and msSIRT2-transgenic mice injected with either vehicle or dex. Sections were stained using WGA, H and E, and Trichrome staining to depict muscle fiber cross sectional area, morphology, and fibrosis, respectively. Scale bars = 50, 75, and 75 µm, respectively. **(I)** Quantification of WGA images represented in Figure 4(H), showing muscle fiber cross sectional area of gastrocnemius muscles in non-transgenic and msSIRT2-transgenic mice administered with either vehicle or dex. Data are shown as median with 25 and 75 percentile. n=3-4, N =702-839. **(J)** Quantification of WGA images represented in Figure 4(H), showing the frequency distribution of cross sectional fiber area of gastrocnemius muscles in non-transgenic and msSIRT2-transgenic mice administered with either vehicle or dex. Data are shown as mean ± s.d. n=3-4. **(K)** Quantification of Masson’s Trichrome images represented in Figure 4(H), showing fibrosis score of gastrocnemius muscles in non-transgenic and msSIRT2-transgenic mice administered with either vehicle or dex. Scoring was done in a blind fashion. Data are shown as median with 25 and 75 percentile. n=5. **(L)** Relative expression of MuRF-1 in the gastrocnemius muscles of msSIRT2-transgenic mice under the vehicle or dex treatment, normalized with the vehicle-treated non-transgenic control. Data are shown as mean ± s.d. n = 5-6. **(M)** Relative expression of Atrogin-1 in the gastrocnemius muscles of msSIRT2-transgenic mice under the vehicle or dex treatment, normalized with the vehicle-treated non-transgenic control. Data are shown as mean ± s.d. n = 5-6. **(N)** Quantification of time the msSIRT2-transgenic and non-transgenic mice endured in the rotating rod after 15 days of administration of either vehicle or dex (10mg/kg/day). Data are shown as mean ± s.d. n = 5-7. The two-tailed unpaired student’s t-test with Welch’s correction (A) Two-tailed Mann-Whitney statistical test (D) Kruskal-Wallis test with Dunn’s multiple comparisons was used for statistical analysis (E, F, G, I, K, L, M, N)

Interestingly, msSIRT2-Tg mice demonstrated improved muscle endurance during the rotarod muscle performance test compared to N-Tg mice under basal conditions, suggesting that SIRT2 overexpression in skeletal muscles may improve muscle function under basal conditions (**Figure 4D**). Following this, we characterized the effect of dex treatment on these mice and found that the msSIRT2-Tg mice were protected from loss in body weight, while N-Tg mice showed a significant reduction in body weight upon dex treatment (**Figure 4E, S6I)**. Similarly, while dex-treated N-Tg mice displayed muscle weight loss in type II muscle fibers, the msSIRT2-Tg mice were protected from it (**Figure 4F, G, S6J, K).** In contrast, the mass of other muscles remained unchanged **(Figure S6L-Q)**. There was also a significant reduction in muscle fiber cross-sectional area in dex-treated N-Tg mice but not in the msSIRT2-transgenic mice (**Figure 4H, I**). Similarly, the frequency of small myofiber bundles was increased significantly upon dex treatment in N-Tg mice but not in the msSIRT2-Tg mice (**Figure 4J**). Moreover, the dex-treated msSIRT2-Tg mice muscle also displayed reduced fibrosis compared to the N-Tg dex-treated controls (**Figure 4H, K**). Furthermore, upon dex treatment, there was an increase in the atrogene expression in both msSIRT2-Tg and N-Tg mice. However, the expression of atrogenes in dex-treated msSIRT2-Tg mice was significantly lower than that of N-Tg mice (**Figure 4L, M**). Interestingly, dex-treated msSIRT2-Tg mice demonstrated significantly better endurance compared to their N-Tg counterparts in rotarod running experiment (**Figure 4N, S6R)**. We then looked at the muscle fiber distribution in the gastrocnemius of these animals and found no significant difference in the muscle fiber types upon dex treatment **(Figure S7A-F**) suggesting that SIRT2 overexpression may protect against dex-induced reduction in muscle endurance and improve overall muscle function in mice. These results indicate that SIRT2 overexpression in skeletal muscles may protect mice from dex-induced muscle atrophy and enhance skeletal muscle performance in mice.

### SIRT2 deficiency disrupts GR signalling pathways involved in muscle homeostasis

Next, to elucidate the mechanism underlying SIRT2-mediated protection from dex-induced atrophy, we characterized the metabolic state of the skeletal muscles. For that, we first examined the mitochondrial COX and SDH staining in the gastrocnemius muscles of SIRT2^-/-^ mice under dex injection (**Figure 5A**). We found no significant difference in the SIRT2^-/-^ muscles with or without dex administration compared to the wild-type mice (**Figure 5B, C**). We then performed the same experiment with msSIRT2-Tg mice and found similar results (**Figure 5D-F**). Further, we assessed oxidative phosphorylation (OXPHOS) and glycolysis in WT and SIRT2^-/-^ primary myocytes and SIRT2 inhibited (AGK2 treated) primary myocytes using a Seahorse XF Extracellular Flux Analyzer by measuring oxygen consumption rate (OCR) and extracellular acidification rate (ECAR), respectively. Depletion of SIRT2 did not alter either OCR or ECAR (**Figure 5G, H, S8A, B)**. Thus, under the conditions tested, the effects of dexamethasone and the contribution of SIRT2 to skeletal muscles appear to be independent of global bioenergetic flux (basal mitochondrial respiration and glycolysis).

**Figure 5.**
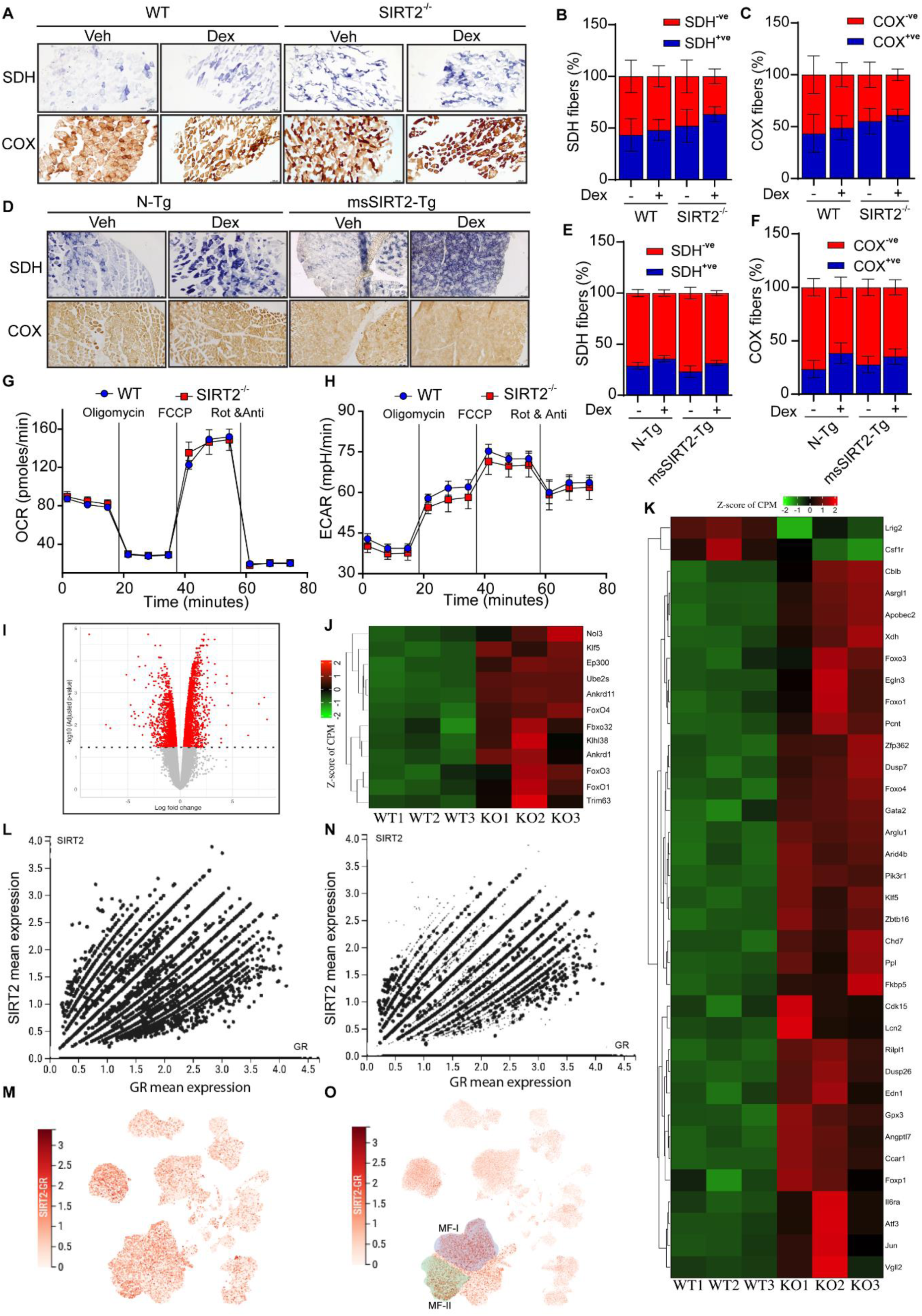
SIRT2 deficiency disrupts GR signalling pathways involved in muscle homeostasis. **(A)** Representative immunohistochemical images showing Succinate dehydrogenase (SDH) and cytochrome oxidase (COX) stain in the gastrocnemius sections of wild-type and SIRT2^-/-^ mice administered with either vehicle or dex (10mg/kg/day). Scale bar = 100 µm. **(B)** Quantification of the percentage of myofibers positive for the SDH staining as shown in Figure 5(A). Data represented as mean ± s.d., n = 3-4. **(C)** Quantification of the percentage of myofibers positive for the COX staining as shown in Figure 5(A). Data represented as mean ± s.d., n = 3-4. **(D)** Representative immunohistochemical images showing Succinate dehydrogenase (SDH) and cytochrome oxidase (COX) stain in the gastrocnemius sections of non-transgenic control and muscle specific SIRT2 transgenic mice administered with either vehicle or dex (10mg/kg/day). Scale bar = 100 µm. **(E)** Quantification of the percentage of myofibers positive for the SDH staining as shown in Figure 5(D). Data represented as mean ± s.d., n = 3-4. **(F)** Quantification of the percentage of myofibers positive for the COX staining as shown in Figure 5(D). Data represented as mean ± s.d., n = 3-4. **(G)** OXPHOS-associated Oxygen Consumption Rate (OCR) in the wild type and SIRT2^-/-^ myotubes measured by mitostress test. Data represented as mean ± s.d., *N*=3, *n* = 3. **(H)** Extracellular Acidification Rate (ECAR) associated with glycolysis in the wild type and SIRT2^-/-^ myotubes, detected by mitostress test. Data represented as mean ± s.d., *N*=3, *n* = 3. **(I)** Representative volcano plot of differentially expressed genes in Gas muscles of wild type and SIRT2^-/-^ mice. 3783 genes are differentially expressed, 2176 are upregulated and 1607 are downregulated in SIRT2 KO mice when compared with wild-type mice. **(J)** Heatmap depicting the GR-interacting GO enriched differentially expressed genes in Gas muscles of wild type and SIRT2^-/-^ mice. **(K)** Heatmap depicting the GR associated differentially expressed genes in Gas muscles of wild type and SIRT2^-/-^ mice. **(L)** Representative graph showing mean expression of GR with respect to SIRT2 in the single cells and single nucleus of various cell types in the young and old individuals. The figures were obtained from muscle aging cell atlas. **(M)** Uniform Manifold Approximation and Projection of SIRT2 and GR. The figures were obtained from muscle aging cell atlas. The figures were obtained from muscle aging cell atlas. **(N)** Representative graph showing mean expression of GR with respect to SIRT2 in the single cells and single nucleus of myofibers in the young and old individuals. **(O)** Uniform Manifold Approximation and Projection of SIRT2 and GR in the myofibers. The figures were obtained from muscle aging cell atlas Two-way ANOVA with Tukey’s multiple comparisons was used for assessing SDH+ve and COX+ve cells (B, C, E, F).

Since we did not find any metabolic change, we then looked for the transcriptome profile in the gastrocnemius muscles of SIRT2-deficient mice. The transcriptome data show that 3,783 genes in the SIRT2-deficient mice were differentially regulated (**Figure 5I**). Our transcriptomic analysis revealed some of the major genes responsible for skeletal muscle atrophy to be highly upregulated **(Figure S8C**). Interestingly, GO analysis also revealed many genes associated with Glucocorticoid Receptor (GR) signalling to be highly upregulated (**Figure 5J**). We further compared our data to that of a recently published research article [44]. This comparative analysis revealed many more genes of GR signalling to be upregulated in the gastrocnemius muscles of SIRT2^-/-^ mice (**Figure 5K**). We also assessed the expression levels of SIRT2 and GR in a previously published dataset in mixed-type intercostal muscles of humans [45]. Our analysis from this single-cell and single nuclei sequencing data shows that upon ageing, reduction in SIRT2 expression correlates with increase in GR expression (**Figure 5L-M**). This correlation was more pronounced in Muscle fiber type I and muscle fibre type II cell population compared to overall cell types sequenced in the study (**Figure 5N-O**).

The glucocorticoid receptor is the primary receptor for the stress hormone cortisol. As an adaptive response to stress, cortisol is known to bind to and activate GR to trigger several downstream signalling pathways that ultimately result in atrophy [18]. Thus, our findings led us to hypothesize that SIRT2 may regulate GR signalling in stress-induced skeletal muscle atrophy. Interestingly, we observed that serum cortisol levels are significantly high in SIRT2^-/-^ mice compared to the wild-type controls (**Figure 6A**).

**Figure 6.**
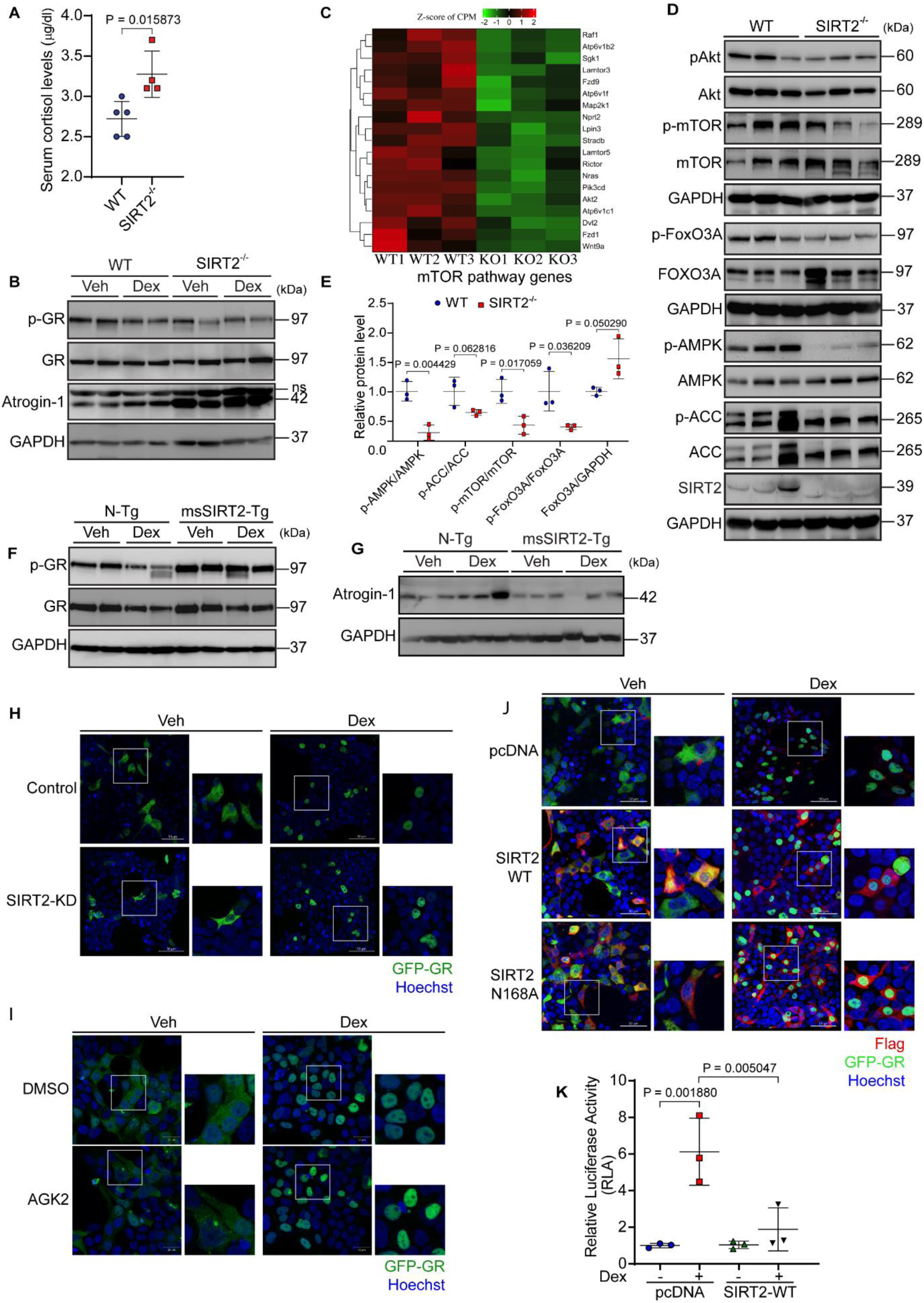
SIRT2 deficiency induces GR signalling in muscle. **(A)** Serum cortisol levels of wild-type and SIRT2^-/-^ mice. Data are shown as mean ± s.d., n = 4-5. **(B)** Representative immunoblot images depicting Atrogin-1 levels and GR levels in the gastrocnemius muscles of SIRT2^-/-^ mice injected with either vehicle or dex. Indicated band sizes are in kilodaltons (kDa). **(C)** Heatmap depicting significantly downregulated genes in SIRT2^-/-^ gastrocnemius muscles that are also involved in MTOR signalling. Upregulated and downregulated genes are represented in red and green, respectively. **(D)** Representative immunoblot images depicting AMPK, mTOR and FoxO3A signaling, in the gastrocnemius muscles of SIRT2^-/-^ mice. Indicated band sizes are in kilodaltons (kDa). **(E)** Scatterplot representing quantification of immunoblots from Figure 6(D). Graph represents relative protein levels. Data are shown as mean ± s.d. n = 3. **(F)** Representative immunoblot images depicting GR signaling in the gastrocnemius muscles of msSIRT2-transgenic mice upon dex injection. Indicated band sizes are in kilodaltons (kDa). **(G)** Representative immunoblot images depicting Atrogin-1 levels in the gastrocnemius muscles of msSIRT2-transgenic mice upon dex injection. Indicated band sizes are in kilodaltons (kDa). **(H)** Representative immunofluorescence images showing GR localization of the vehicle and dex-treated HEK293T knocked down with SIRT2. Cells were overexpressed with GFP-GR (green) and then stained with Hoechst (blue). GR intensity has been modified to represent its localization better. Scale bar = 50 µm. **(I)** Representative immunofluorescence images showing GR localization upon AGK2 treatment of the vehicle and dex-treated HEK293T. Cells were overexpressed with GFP-GR (green) and then stained with Hoechst (blue). GR intensity has been modified to represent its localization better. Scale bar = 50 µm. **(J)** Representative immunofluorescence images showing GR localization of the vehicle and dex-treated HEK293T overexpressed with SIRT2 WT and its mutant (SIRT2 N168A). Cells were overexpressed with GFP-GR (green) and then stained with Flag (red), and Hoechst (blue). GR and flag intensity has been modified to represent its localization better. Scale bar = 50 µm. **(K)** Relative luciferase activity depicting GR’s activity as a transcription factor. HEK293T cells were transfected with MMTV-Luc plasmid with either pcDNA or SIRT2-wild-type plasmids under the vehicle and dex-treated conditions. pRL-CMV was used as control. Data are shown as mean ± s.d. n = 3. Two-tailed Mann-Whitney statistical test (A) Multiple t tests with Holm-Sidak method to correct for multiple comparision (E) Two-way ANOVA with Tukey’s multiple comparisons test was used for statistical analysis (K).

Since muscle atrophy is caused by loss of protein homeostasis, we hypothesized that SIRT2^-/-^ muscles may display glucocorticoid-mediated deregulation of protein synthesis and degradation, leading to muscle atrophy. To evaluate the role of SIRT2 in regulating GR activity in atrophy-associated loss of proteostasis, we first assessed changes in the activity of GR in the skeletal muscles of SIRT2^-/-^ mice. Previous reports indicate that inhibitory phosphorylation of GR at S226 reduces its activity [46]. Thus, we assessed the phosphorylation of GR as an indicator of its activity. We found that dex-injected wild-type mice displayed a marked reduction in the inhibitory phosphorylation of the glucocorticoid receptor (S226), indicating increased glucocorticoid receptor activity upon dex injection (**Figure 6B**). Similar to dex injection in wild-type mice, the phosphorylation of GR was also downregulated in the gastrocnemius of SIRT2^-/-^ mice with or without dex, indicating increased GR activity (**Figure 6B**). Concomitant with decreased phosphorylation of GR, the gastrocnemius muscles of SIRT2^-/-^ mice showed increased levels of Atrogin-1 in the vehicle and dex-treated conditions (**Figure 6B**). Next, to understand how SIRT2 may regulate GR phosphorylation, we assessed the expression of Protein Phosphatase 2A (PP2A). It is known that PP2A dephosphorylates GR and increases its nuclear localization [47]. We observed that both RNA and protein levels of PP2A were highly upregulated in SIRT2-deficient mice treated with dex **(FigureS8D-F)**. We speculate that depleting the levels of SIRT2 may indirectly increase the activity of PP2A. These data indicate that SIRT2 deficiency may activate GR signalling to induce muscle atrophy.

To delineate the molecular mechanism underlying stress-induced GR-associated remodelling of muscle protein homeostasis in SIRT2^-/-^ mice, we assessed the protein levels of key molecules involved in the the regulation of protein synthesis, including Akt and its downstream targets like mTOR and FoxO3A, AMPK and acetyl CoA carboxylase (ACC). Under chronic stress conditions, activated GR inhibits the IGF1-PI3K-AKT pathway responsible for protein synthesis and muscle growth [18]. Subsequently, FOXO activation upregulates several atrophy-related genes, including MuRF-1 and Atrogin-1 [48–52]. GO analysis of RNA sequencing data derived from SIRT2^-/-^ muscle indicated downregulation of genes involved in the mTOR signalling pathway (**Figure 6C**). Consistent with this, we found a significant reduction in Akt (S473) phosphorylation as well as mTOR (S2448) phosphorylation in SIRT2^-/-^ muscle tissue (**Figure 6D**). We also observed reduced protein synthesis in SIRT2^-/-^ mice, and this phenotype was aggravated upon dex treatment **(Figure S8G-J)**, suggesting that SIRT2^-/-^ muscles may display glucocorticoid-mediated deregulation of protein synthesis and degradation, leading to muscle atrophy. Since Akt-mediated mTOR phosphorylation was reduced, we checked for a critical Akt target, FoxO3A, which serves as an important transcription factor of Atrogin-1 and MuRF1 [53, 54]. Previously reports have shown that FoxO3A is an early target of GR, which is transcriptionally upregulated upon GR activation [55, 56]. We observed a reduction in inhibitory phosphorylation of FoxO3A and an increase in FoxO3A expression, indicating increased activation of FoxO3A (**Figure 6D, E**). Previous studies have also shown that under chronic stress conditions, AMPK modulates protein synthesis and degradation in muscle atrophy [57–59]. We found that phosphorylation of AMPK and the phosphorylation on its downstream target ACC, were both reduced in SIRT2^-/-^ muscle (**Figure 6D, E**), suggesting that GR might regulate protein synthesis in SIRT2^-/-^ muscle independent of AMPK. Altogether, these results suggest that SIRT2 depletion may inhibit protein synthesis and promote protein degradation, thereby inducing muscle atrophy in mice.

Since our *in vitro* results indicate that SIRT2 overexpression may ameliorate glucocorticoid-mediated muscle atrophy, we hypothesized that SIRT2 overexpression might reduce GR expression and activity in dex-injected mice. As expected, the msSIRT2-Tg mice showed increased GR phosphorylation under the vehicle or dex-injected conditions (**Figure 6F**). In addition to increased GR phosphorylation, we also observed reduced levels of Atrogin-1 in the gastrocnemius muscles of msSIRT2-Tg mice either injected with vehicle or dex (**Figure 6G**). Furthermore, we observed that msSIRT2-Tg mice were protected against dex-induced inhibition of protein synthesis (**Figure S8K, L)**.

Next, we wanted to understand whether SIRT2 levels affect GR nuclear localization. To assess this, we over-expressed GFP-GR in SIRT2-depleted cells and observed GR localization under control and dex-treated conditions. We observed no significant change in GR localization under dex treatment (**Figure 6H**). Consistent with this, GR localization also remained unaffected in AGK2-treated cells (**Figure 6I**). Interestingly, GR localization was unaffected even in cell populations overexpressing SIRT2 wild-type and catalytically inactive SIRT2 mutant (N168A) proteins under dex-treated conditions (**Figure 6J**). However, we found that wild-type SIRT2 overexpression reduced GR activity *in vitro* (**Figure 6K**). Since activated GR is known to induce transcription of atrophy related genes directly by binding to their promoter region [52, 56, 60], we performed Chromatin immunoprecipitation assay (ChIP) in muscles of WT and SIRT2^-/-^ mice. We observed increased DNA-binding of GR under SIRT2 deficiency. Specifically, GR occupancy was increased at GR target promoters (FKBP5, TAT, Slc3a1 intron) as well as FOXO3A target promoters (Atrogin-1 and MuRF-1) in SIRT2^-/-^ muscles compared to its WT counterparts **(Figure S9 A-F)**.Together, these results suggest that SIRT2 might play a critical role in muscle atrophy by catalytically regulating GR activity under stress conditions.

### SIRT2 binds to the glucocorticoid receptor to regulate muscle atrophy

Previous reports indicate that acetylation of GR regulates its activity [61]. Since SIRT2 is a known deacetylase, we speculated that SIRT2 may bind to and de-acetylate GR to regulate its activity. To evaluate whether SIRT2 may directly interact with and deacetylate GR, we over-expressed SIRT2 and GR *in vitro* and performed co-immunoprecipitation. We observed that SIRT2 interacts with GR under basal conditions (**Figure 7A, S10A).** Previous studies have reported that SIRT2 regulates GR activity via Hsp90, by deacetylating HSP90 in neuronal cells [62]. Thus, to understand if SIRT2 may regulate GR in Hsp90-dependent manner in skeletal muscles, we first assessed Hsp90 expression in SIRT2-deficient muscles treated with dex. We found an increase in Hsp90 mRNA as well as protein levels upon dex treatment in both WT and SIRT2^-/-^ mice (**Figure S10B-D**). Since Hsp90 is known to be required for GR maturation and stability, increase in Hsp90 expression may be attributed to increased GR stability. However, to understand if Hsp90 is involved in SIRT2-mediated GR regulation in skeletal muscles, we immunoprecipitated GR in muscle tissue derived from SIRT2^-/-^ mice. We found an increase in Hsp90 and GR binding in SIRT2^-/-^ skeletal muscles compared to the control (**Figure 7B**). However, interestingly, we found no significant difference in Hsp90 acetylation compared to control (**Figure 7B**). These results suggest, although Hsp90 may be required for GR stability, it is not a deacetylation target of SIRT2 and may not be involved in SIRT2-GR binding and regulation in skeletal muscles.

**Figure 7.**
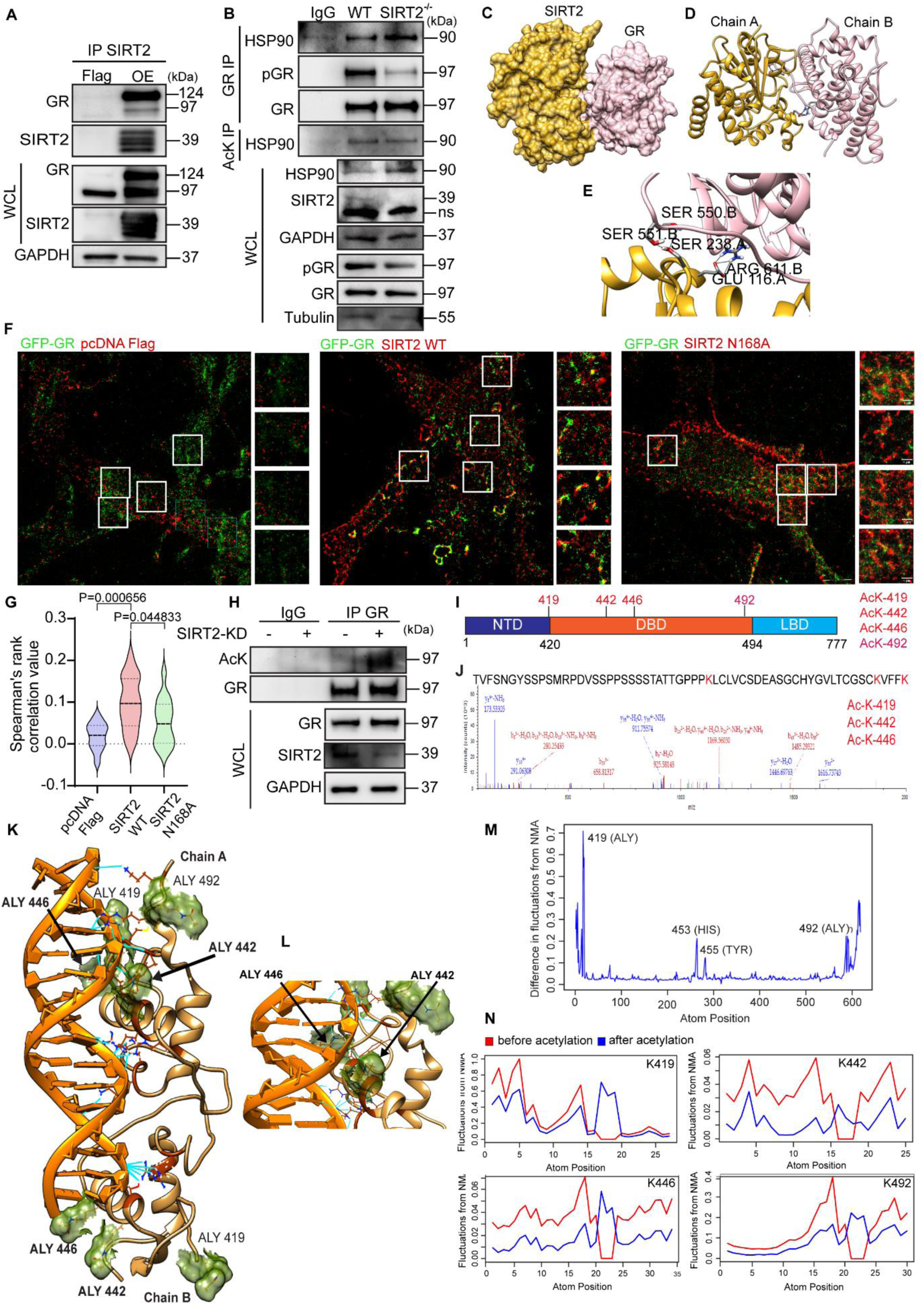
SIRT2 interacts with the glucocorticoid receptor and regulates GR signalling. **(A)** Representative immunoblot images showing the interaction of SIRT2 with GR. SIRT2 and GR were overexpressed in HEK293T cells, and SIRT2 was immunoprecipitated and probed for GR. Whole cell lysate (WCL) was also immunoblotted to check the loading. Indicated band sizes are in kilodaltons (kDa). **(B)** Representative immunoblot images showing the interaction of HSP90 with GR and acetylation of HSP90. GR and acetyl lysine was immunoprecipitated and probed for HSP90. Whole cell lysate (WCL) was also immunoblotted to check the loading. Indicated band sizes are in kilodaltons (kDa). **(C)** Surface representation of the docked crystal structure of SIRT2 monomer (PDB code: 5Y0Z, coloured in golden) and Ligand binding domain (LBD) crystal structure of Glucocorticoid Receptor (GR, PDB code: 7PRX, coloured in pink). **(D)** Cartoon representation of the SIRT2-GR docked complex. Chain A represents the SIRT2 monomer, and chain B in the docked complex represents the LBD of GR. The black lines indicate hydrogen bonds formed between the two proteins. **(E)** Zoomed-in representation of the residues of SIRT2 and GR involved in hydrogen bonding. **(F)** Representative color aligned STORM microscopy images after chromatic aberration correction, showing the interaction of SIRT2 with GR in C6 glioma cells overexpressing flag-pcDNA, flag-SIRT2-WT and flag-SIRT2-N168A mutant with GFP-GR. Representative 250*250 pixel sized cropped regions on which colocalization analysis was done. Yellow regions show co-localization of single SIRT2 and single GR molecules. Scale bar=5um (whole cell images), 1um (cropped regions). **(G)** Quantification of figure 7(F) depicted in the form of Spearman’s rank correlation value. **(H)** Representative immunoblot images showing the GR as an acetylated protein. SIRT2 was knocked down, and GR was overexpressed in C2C12 cells, and GR was immunoprecipitated and probed for pan-acetylated lysine. Whole cell lysate (WCL) was also immunoblotted to check the loading. Indicated band sizes are in kilodaltons (kDa). **(I)** Schematic diagram showing acetylated lysine residues in the GR under SIRT2 inhibited cells. The GR is divided into the N-terminal domain (NTD), the DNA-binding domain (DBD), and the Ligand binding domain (LBD). Four Lysine residues that are acetylated during SIRT2-inhibited conditions are displayed. The three novel residues (K419, 442, 446) are shown in red. **(J)** Graph depicting the mass-to-charge ratio of a peptide from GR, showing acetylation of two lysine residues (K442, 446) **(K)** Structure of human glucocorticoid receptor complexed to CCL2 NF-kB response element [65] with 7 acetylated lysine residues (ALY), 4 in chain A (419, 442, 446, 492) and 3 in chain B (419, 442, 446). The DNA fragment is colored orange and the protein dimer in light brown. Interacting interfacial residues are represented in brown and the olive surfaces mark the acetylated lysine residues. Hydrogen bonds formed between the protein and DNA are shown in cyan. **(L)** Zoomed-in view of two acetylated lysine residues (442 and 446) that interact with DNA. **(M)** Difference in fluctuations, obtained from all-atom based ENM-NMA for all atoms in all residues of one chain of the glucocorticoid protein dimer. The residues showing high difference in their fluctuation profile are marked. **(N)** Standardized square fluctuations, obtained from all-atom-based ENM-NMA for acetylated residues in one chain of the GR dimer, before and after acetylation. Ordinary one-way ANOVA with Tukey’s multiple comparisons test was used for statistical analysis (G).

To further characterize the interaction between SIRT2 and GR, we performed in silico analysis. To predict the binding interface between SIRT2 and GR, we performed protein-protein docking using HADDOCK (High Ambiguity Driven protein-protein DOCKing), a widely used information-driven flexible docking approach for modeling biomolecular complexes. As there is no resolved crystal structure of full-length GR, we initially examined the AlphaFold-predicted GR structure. However, the predicted structure was largely in an open conformation, with the majority of the backbone displaying low to very low per-residue confidence scores (pLDDT < 70), particularly in flexible regions **(Figure S10E).** This structural uncertainty makes the AlphaFold predicted model unreliable for docking, as poorly defined regions can introduce significant inaccuracies in interface prediction. Given these challenges, we opted for a docking approach using HADDOCK, which leverages experimentally derived or predicted interaction restraints to guide complex formation. The docking was restricted to the ligand-binding domain (LBD) of GR, as the full-length structure - including the N-terminal domain, DNA-binding domain (DBD), and LBD - is not resolved and unavailable in the RCSB Protein Data Bank. While SIRT2 deacetylates residues in the DBD, this does not necessarily imply direct docking to this domain. The preference for LBD docking is based on the fact that post-translational modifiers, including deacetylases, interact with nuclear receptors primarily through regulatory domains such as the LBD rather than directly interacting with DNA-binding regions [63–65]. In this context, SIRT2 may engage with the LBD, potentially influencing the DBD through allosteric effects or transient interactions. The docking solutions were ranked using the HADDOCK scoring function, which integrates van der Waals, electrostatic, desolvation, and distance restraint energies. The top-ranked cluster, with the most favourable HADDOCK score (−83.2), was selected, with energy contributions as follows: van der Waals energy (−37.8 kcal/mol), electrostatic energy (−201.6 kcal/mol), and desolvation energy (−5.1 kcal/mol). The analysis of the SIRT2-GR complex revealed five hydrogen bonds at the interaction interface, involving specific residues on both proteins (**Table 1**). While hydrogen bonding is not the primary driver of protein-protein interactions compared to hydrophobic forces, shape complementarity, and overall energetic contributions such as van der Waals and electrostatic interactions, it remains an important factor in molecular recognition and specificity. In in-silico docking studies, hydrogen bonds serve as a proxy for potential interaction sites, aiding in the structural prediction of docked complexes. HADDOCK evaluates these interactions through an energy-based scoring function that integrates multiple energetic terms to identify the most favourable binding conformations. These findings suggest that these interactions may contribute to the stability and specificity of the SIRT2-GR complex formation (**Figure 7C-E**).

**Table 1.** Specific residues involved in H-bonds in SIRT2-GR complex.

| Donor (D) | Acceptor (A) | D..A distance<br>(Å) |
| --- | --- | --- |

| Residue | Position | Atom | Residue | Position | Atom |  |
| --- | --- | --- | --- | --- | --- | --- |
| SER | 238.A | OG | SER | 551.B | OG | 2.776 |
| SER | 550.B | N | SER | 238.A | OG | 2.833 |
| ARG | 611.B | NH1 | GLU | 116.A | OE1 | 2.632 |
| ARG | 611.B | NH2 | GLU | 116.A | OE1 | 3.299 |
| ARG | 611.B | NH2 | GLU | 116.A | OE2 | 3.377 |

Next, to experimentally verify SIRT2-GR interaction, we performed an immunofluorescence-based colocalization assay, indicating that SIRT2 binds to GR under basal conditions. Using thresholded Mander’s coefficients (tM1 and tM2) and Spearman’s rank correlation, we noted an increase in co-localization between GR and SIRT2 upon overexpression of Flag-SIRT2 compared to pcDNA control (**Figure S10F-I)**. To fortify these findings, we assessed the proximity of SIRT2 and GR using direct Stochastic Optical Reconstruction Microscopy (dSTORM), which enables nanoscale interaction between the interacting proteins with subpixel resolution. Following dSTORM, quantification with Mander’s and Spearman’s rank correlation values revealed positive correlation and colocalization between SIRT2 and GR. The results indicate that the interaction between SIRT2 and GR was not significantly altered when a catalytically inactive mutant of SIRT2 (N168A) was used instead of SIRT2 WT (**Figure 7F, G, S10J, K)**. GR has a molecular weight of approximately 97 kDa and is a globular protein with a dimension of about ∼8-10 nm, while SIRT2 has a molecular weight of approximately 43 kDa and a dimension of ∼ 5-7 nm. The localization precision of dSTORM after immunolabeling (∼20 nm), is within the same order of magnitude as the combined approximate dimensions of GR and SIRT2, making it unlikely that an additional protein consistently bridges the observed proximity. Thus, the colocalization of proteins observed are indicative of direct or very close interaction. These quantitative metrics suggests that SIRT2 and GR share overlapping spatial distributions in the cytoplasm, confirming direct association of SIRT2 with GR.

Since SIRT2 is known to modulate protein functions via deacetylation, we investigated whether GR acetylation is altered upon SIRT2 depletion. We observed that GR is increasingly acetylated under SIRT2-depleted conditions *in vitro* (**Figure 7H**). These data suggest that SIRT2 may directly deacetylate GR to modulate its activity. To further identify the residues at which the SIRT2 may deacetylate the glucocorticoid receptor, we performed mass spectrometry analysis using the glucocorticoid receptor protein isolated from GR overexpressing cells treated with SIRT2 inhibitor AGK2. We found that four lysine residues in and around the DNA binding domain of GR are acetylated (K419, 442, 446, 492) under AGK2-treated conditions (**Figure 7I, J**). We then adopted an *in silico-*based approach to understand how acetylation at these residues may impact GR activity. The crystal structure of the GR DNA-binding domain complexed to the CCL2 NF-kB response element does not have any lysine residues of interest acetylated. Hence, we performed in-silico acetylation on the SIRT2-specific lysine residues (419, 442, 446, 492 of chain A and 419, 442, 446 of chain B), followed by energy minimization of the acetylated model. The acetylated GR protein complexed to genomic NF-kB response element is shown in (**Figure 7K, L**). All the acetylated lysine residues are shown in olive and surface representation. The interfacial residues that interact with DNA are shown as brown-coloured heteroatoms. The H-bonds formed between the protein and DNA are represented in cyan. The total interaction energy of the acetylated protein-DNA complex is −1.4010765e+06 KJ/mol. This is substantially lower than the total energy of interaction before acetylation, which was found to be −1.3899310e+06 KJ/mol. This substantial reduction in interaction energy after acetylation can be associated with better DNA-binding ability of the acetylated protein than its unacetylated counterpart. Incidentally, three additional H-bonds exist between the protein and DNA in the acetylated form compared to the unacetylated form. **Table 2** lists the H-bonds observed between the acetylated protein and the DNA. This also includes all the other hydrogen bonds noted in the unacetylated form. The three additional H-bonds are highlighted in blue. Out of all the acetylated lysine residues, only two (442, 446) were found to be interacting with DNA.

**Table 2.** H-bonds between the acetylated GR protein and the DNA.

| Donor (D) |  |  | Acceptor (A) |  |  | D..A distance (Å) | D-H..A distance (Å) |
| --- | --- | --- | --- | --- | --- | --- | --- |
| Residue | Position | Atom | Residue | Position | Atom |  |  |
| CYS | 431.A | N | DG | 4.C | O2P | 2.931 | 1.936 |
| TYR | 433.A | N | DG | 5.C | O2P | 2.698 | 1.693 |
| ALY | 442.A | NZ | DG | 5.C | N7 | 3.37 | 2.468 |
| ALY | 446.A | NZ | DA | 6.C | O1P | 2.857 | 1.867 |
| ARG | 447.A | NH1 | DA | 8.D | N7 | 3.424 | 2.573 |
| ARG | 447.A | NH2 | DA | 7.D | O2P | 2.675 | 1.672 |
| ARG | 470.A | NE | DA | 8.D | O1P | 2.704 | 1.714 |
| ARG | 470.A | NH2 | DA | 8.D | O5' | 2.958 | 2.121 |
| ARG | 470.A | NH2 | DT | 9.D | O2P | 2.694 | 1.7 |
| LYS | 471.A | NZ | DC | 13.C | O1P | 2.7 | 1.647 |
| ARG | 477.A | NH1 | DA | 8.D | O2P | 2.668 | 1.715 |
| ARG | 477.A | NH2 | DA | 8.D | O2P | 2.765 | 1.849 |
| ARG | 491.A | NH1 | DG | 4.C | N3 | 2.978 | 2.046 |
| ARG | 491.A | NH1 | DG | 5.C | O4' | 2.857 | 1.955 |
| ARG | 491.A | NH2 | DG | 4.C | N3 | 2.983 | 2.043 |
| LYS | 494.A | NZ | DT | 16.D | O1P | 2.665 | 1.618 |
| SER | 440.B | OG | DC | 15.C | O2P | 2.93 | 2.233 |
| SER | 440.B | OG | DC | 15.C | O5' | 2.972 | 2.062 |
| ARG | 470.B | NE | DC | 15.C | O1P | 2.704 | 1.73 |
| ARG | 470.B | NH2 | DC | 15.C | O1P | 2.937 | 2.048 |
| ARG | 477.B | NH1 | DC | 15.C | O2P | 2.94 | 2.094 |
| ARG | 477.B | NH2 | DC | 15.C | O2P | 2.67 | 1.684 |

Further, we performed an all-atom Normal Mode Analysis (NMA) for both the unacetylated and acetylated form of the GR Protein-DNA complex. We observed that, in general, the flexibility of GR decreases in the acetylated form as compared to the unacetylated form (**Figure 7M**), indicating the difference in flexibility between all residues of the acetylated and unacetylated form in one of the dimer chains, suggesting lower flexibility of GR in the acetylated form compared to the unacetylated form. The acetylated residues, particularly showing lower flexibility after acetylation, are shown in (**Figure 7N**). The reduction in flexibility after acetylation indicates a tighter interaction between the GR protein and DNA. This is consistent with the total interaction energy calculation and the formation of additional interactions. Thus, our structural and flexibility analysis indicates that upon acetylation, the DNA-protein interactions get enhanced, and the flexibility associated with the binding site within the protein is reduced (**Figure 7K, L**).

### SIRT2 deacetylates critical GR lysine residues to regulate GR signalling

We experimentally validated these findings by generating a charge mimic for acetylation (K→Q) and deacetylation (K→R) of GR at 442 and 446 residues. To our surprise, we found that (K→R, K→Q) mutations at K442 and K446 disrupted the DNA binding activity of the endogenous glucocorticoid receptor (**Figure 8A**). We speculate that since the mutations made were charge mimics and not accurate mimics for the acetylation/deacetylation of lysine, we observe a disruption in the GR activity with either mutation. These results suggest that K442 and K446 residues are critical for DNA binding, and any modification at these sites may result in the disruption of the DNA binding activity of the GR.

**Figure 8.**
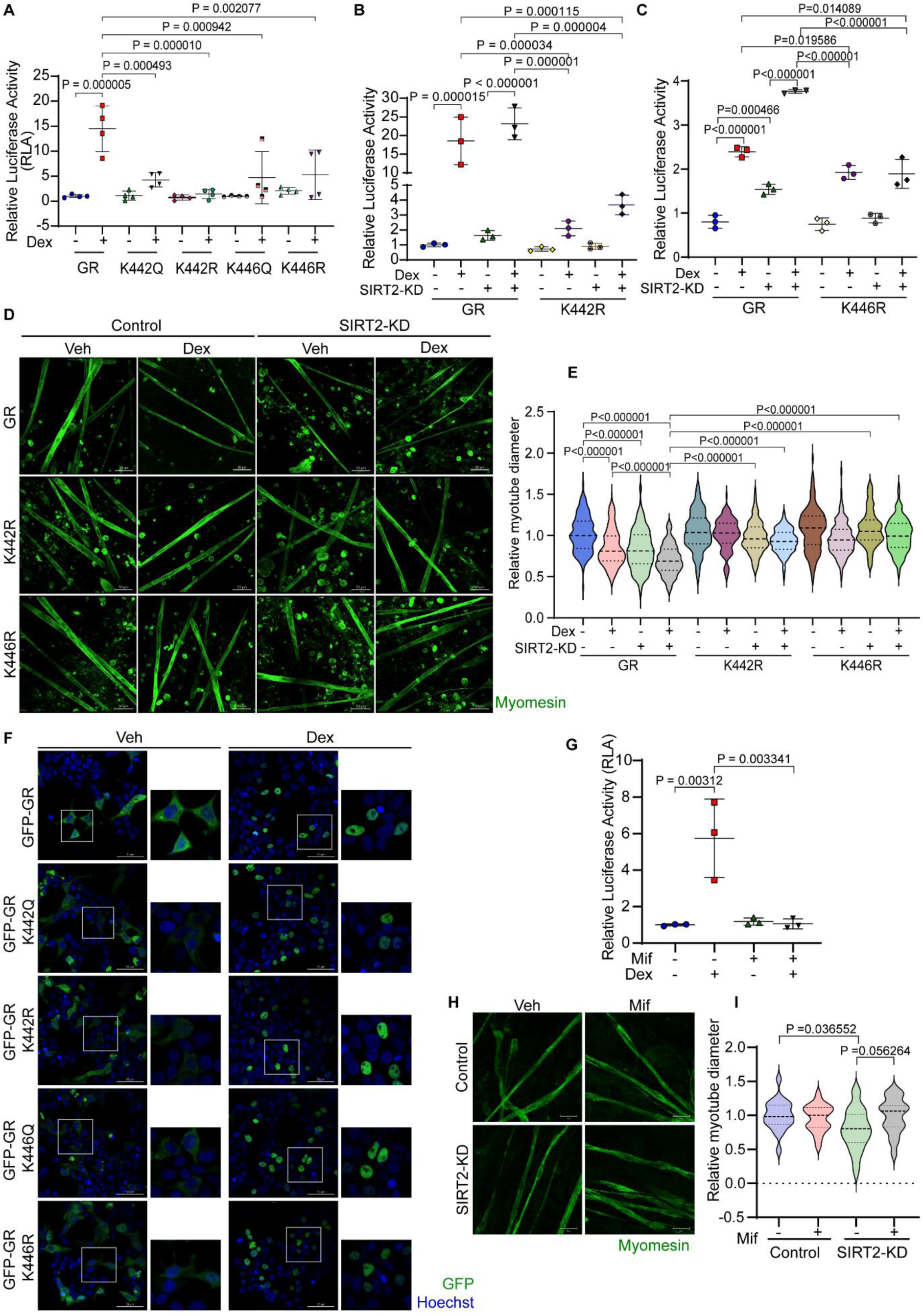
SIRT2 deacetylates critical GR lysine residues to regulate GR signalling. **(A)** Quantification of luciferase activity showing activities (as a transcription factor) of various GR mutants. Wild-type GR along with the mutants where lysine residues K442 and K446 were mutated to either K→Q or K→R and then transfected in HEK 293T cells with MMTV-Luc plasmid. pRL-CMV was used as control. Data are shown as mean ± s.d. n = 4. **(B)** Quantification of luciferase activity showing GR activity (as a transcription factor) in SIRT2 knocked down cells when overexpressed with GFP-GR and GFP-GR K442R. Lysine residue K442 was mutated to K→R and then transfected in HEK 293T cells with MMTV-Luc plasmid. pRL-CMV was used as control. Data are shown as mean ± s.d. n = 3. **(C)** Quantification of luciferase activity showing GR activity (as a transcription factor) in SIRT2 knocked down cells when overexpressed with GFP-GR and GFP-GR K446R. Lysine residue K446 was mutated to K→R and then transfected in HEK 293T cells with MMTV-Luc plasmid. pRL-CMV was used as control. Data are shown as mean ± s.d. n = 3. **(D)** Representative immunofluorescence images showing diameter of the primary rat myotubes in control and SIRT2-depleted conditions along with overexpression of WT GR and its mutants, K442R and K446R when treated with vehicle or dex. Myotubes were stained with antibodies against myomesin (green). Scale bar = 50 µm. **(E)** Quantification of relative primary rat myotube diameters of the representative images shown in Figure 8(D) myotubes in control and SIRT2-depleted conditions along with overexpression of WT GR and its mutants, K442R and K446R when treated with vehicle or dex. Data are shown as median with 25 and 75 percentiles. n=2 **(F)** Representative immunofluorescence images showing GR localization of the vehicle and dex-treated HEK293T under various GR mutations (K442Q, K442R, K446Q, and K446R). Cells were overexpressed with GFP-GR (including mutants) (green) and then stained with Hoechst (blue). GR intensity has been modified to represent its localization better. Scale bar = 50 µm. **(G)** Quantification of luciferase activity of GR as a transcription factor when treated with dex and Mifepristone (Mif). GR was overexpressed in HEK293T cells with MMTV-Luc plasmid. pRL-CMV was used as control. Data are shown as mean ± s.d. n = 3. **(H)** Representative immunofluorescence images displaying the myotubes diameter in control and SIRT2-depleted conditions when treated with vehicle or mifepristone (Mif). Myotubes were stained with antibodies against myomesin (green). Scale bar = 50 µm. **(I)** Quantification of relative myotube diameters of the representative images shown in Figure 8(H) in mif-treated primary myotubes under SIRT2 depleted conditions, normalized with the vehicle-treated control. Data are shown as median with 25 and 75 percentiles. n=3. Ordinary one-way ANOVA with Tukey’s multiple comparisons test (A) Two-way ANOVA with Tukey’s multiple comparisons test (B, C, G, I) Kruskal-Wallis test with Dunn’s multiple comparisons (E) statistical tests were used.

Since we found that K→R mutation suppresses GR activity, we next wanted to understand how this mutation would affect GR activity when SIRT2 levels are downregulated. We found that K442R, and K446R overexpression drastically reduces GR activity, which is irrespective of SIRT2 expression levels (**Figure 8B, C**). We found that overexpressing K442R and K446R rescues the SIRT2 depletion-mediated myotube fiber reduction (**Figure 8D, E**). These data suggest that constitutive deacetylated GR at Lys442 and 446 protects from stress-mediated muscle atrophy. Further, these findings also indicate that SIRT2 may affect muscle atrophy involving this critical residue. Next, to understand whether the SIRT2-dependent deacetylation status of GR affects its nuclear localization, we overexpressed the wild-type GR and GR K442 and GR K446 mutants in cells and found that the mutant GR’s nuclear localization was unaffected in vehicle or dex-treated conditions (**Figure 8F**). Since our results indicate that SIRT2 is responsible for reducing muscle atrophy by inhibiting endogenous GR-DNA binding activity, we performed rescue experiments using mifepristone (Mif), a GR antagonist. We observed rescue of myotube fibre diameter and atrophy-like phenotype in mifepristone treated SIRT2-depleted myotubes (**Figure 8G-I**). These data indicate that SIRT2 ameliorates dex-induced, GR-mediated muscle atrophy. Altogether, the findings in this paper reflect that SIRT2 directly binds to, acetylates, and suppresses GR’s DNA binding activity, thereby protecting against stress-induced skeletal muscle atrophy.

## DISCUSSION

Stress-induced skeletal muscle atrophy is a defining cause of severe discomfort and death in various pathologies, including cardiovascular diseases (CVDs), renal failure, arthritis, cancer, and diabetes [4–9]. In the current study, using *in vitro* and animal model systems, we characterize a novel role for SIRT2 as a critical molecular regulator of GR signalling in dexamethasone-associated stress-induced muscle atrophy. We show that SIRT2 expression is reduced in dex-treated muscles, and depletion of SIRT2 is associated with the development of skeletal muscle atrophy. On the other hand, muscle-specific SIRT2 overexpressing mice are protected from skeletal muscle atrophy. Mechanistically, we demonstrate that SIRT2 directly binds to, deacetylates and negatively regulates GR activity in skeletal muscles by reducing its DNA-binding affinity, without affecting its subcellular localisation when induced with dexamethasone. Altogether, we show that SIRT2 mediates GR signalling in skeletal muscles to protect against stress-induced skeletal muscle atrophy.

In stress-induced skeletal muscle atrophy, activated GR is also known to upregulate the expression of FOXO3a, a transcription factor known to target and enhance atrogene expression [66]. Interestingly, previous reports have also identified FOXO3a as a deacetylation target of SIRT2 in calorie restriction and under oxidative stress [67]. Although not significantly, we observe increased GR occupancy at FOXO3a target promoters in SIRT2^-/-^ muscles. Increased promoter occupancy correlates with the upregulated mRNA levels of atrogenes, as reflected in our RNA-Sequencing dataset and qPCR analysis. Thus, the effect of SIRT2-mediated regulation of muscle atrophy may likely result from the synergistic effect of SIRT2 on both the glucocorticoid receptor and FOXO3a, culminating in resistance to stress-induced atrophy.

GR activation has been associated with reduced muscle protein synthesis, a characteristic feature of muscle atrophy [68, 69]. In our study, we show reduced protein synthesis and enhanced protein degradation in SIRT2^-/-^ mice, indicating that active GR may mediate the effects of chronic stress and induce muscle atrophy expression by disrupting muscle protein homeostasis in SIRT2^-/-^ mice. The reduction in protein synthesis in SIRT2^-/-^ mice may be attributed to changes in critical molecular regulators of protein homeostasis downstream of GR activation, as observed. These include a decrease in p-Akt and p-mTOR levels, suggesting inhibition of mTOR signalling that is critical for protein synthesis. Additionally, FOXO3a, a transcription factor known to increase expression of atrogenes that promote protein degradation, is also upregulated in SIRT2^-/-^ mice. Altogether, a reduction in protein synthesis and an increase in protein degradation result in the loss of muscle protein homeostasis that culminates in skeletal muscle atrophy in SIRT2^-/-^ mice.

Interestingly, SIRT2^-/-^ animals display increased haemoglobin content and PVC/HCT percentage, suggesting improved blood oxygen carrying capacity. To understand if this factor impinges on muscle endurance capacity and homeostasis in SIRT2^-/-^, we evaluated oxidative phosphorylation (OCR) and glycolysis (ECAR). We found no significant differences in basal or maximal respiration, ATP production, or glycolytic rate between WT and SIRT2^-/-^ primary myocytes or in SIRT2-inhibited primary myocytes. We speculate that this, alongwith perturbed protein homeoxostasis downstream of GR activation in SIRT2^-/-^ mice may, at least in part, explain the poor muscle performance observed during the rotarod test with these mice, despite their elevated haemoglobin content and PVC/HCT percentage.

GR is known to undergo several post-translational modifications that regulate its activity. In addition to deacetylation, phosphorylation is one such notable post-translational modification that is known to inhibit GR activity [70]. GR phosphorylation is tightly regulated by an interplay between the GR-inhibitory kinase, JNK and the GR-activating phosphatase, PP2A [47, 71, 72]. We show that pGR levels are reduced in SIRT2^-/-^ mice, indicating increased GR activity upon SIRT2 depletion. This is concomitant with an increase in PP2A expression in these mice. These results suggest that SIRT2 depletion may result in increased PP2A-mediated dephosphorylation and activation of GR, although the molecular mechanism underlying SIRT2-mediated regulation of PP2A remains to be understood. Interestingly, we have previously shown that SIRT2 deacetylates and activates the GR kinase, JNK [73]. Thus, we expect that in SIRT2^-/-^ mice, JNK-mediated phosphorylation of GR may be reduced, potentially accounting for increased GR activity. In this manner, in addition to direct GR deacetylation, SIRT2 may also regulate GR activity indirectly via PP2A and JNK-dependent pathways.

Previously, studies using *in vitro* model systems have shown that SIRT2 indirectly regulates GR via deacetylation of GR-associated Hsp90 chaperone protein in neuronal systems [62]. In skeletal muscles, we do observe an increase in Hsp90 mRNA and protein levels upon dex treatment in both wild-type and SIRT2^-/-^ skeletal muscles. Since Hsp90 is known to be required for GR maturation and stability, an increase in HSP90 expression may increase GR stability upon dex treatment. We also observe an increase in HSP90 and GR binding in SIRT2^-/-^ skeletal muscles. However, contrary to neuronal cells, we do not observe an increase in HSP90 acetylation in SIRT2^-/-^ skeletal muscles when compared to the control. These results suggest that although HSP90 may be required for GR stability, it is not a deacetylation target of SIRT2 in skeletal muscles. These differences in findings indicate distinct tissue-specific mechanisms for SIRT2-mediated regulation of GR in mice.

Activation of SIRT2 regulates autophagy to protect against muscle atrophy [34]. This study primarily used sirtinol, a non-specific Sirtuin inhibitor known to affect SIRT1 and SIRT2. To overcome the limitations of *in vitro* and whole-body transgenic animal model systems and further our understanding of the muscle-specific role of SIRT2 in GR-mediated atrophy, we have developed and characterised a skeletal muscle-specific SIRT2-transgenic mouse model in this study. Interestingly, we observed that muscle-specific SIRT2-transgenic mice demonstrate significantly enhanced muscle endurance during the rotarod muscle performance test compared to control non-transgenic mice. This indicates that SIRT2 may contribute to maintaining muscle function and health and ameliorating stress-induced muscle atrophy.

While our study has several strengths, including the use of genetic models like knock-out and muscle-specific transgenic mice to understand the role of SIRT2 in skeletal muscle atrophy, there are some limitations. Although we have used SIRT2^-/-^ and SIRT2-msTg mice to demonstrate that SIRT2 negatively regulates GR activity to protect against skeletal muscle atrophy, we have not used GR mutant (KR and KQ) knock-in mice to further validate these findings. We have also not performed ChIP-seq or RNA-Seq with KR mutants of GR to validate that acetylation of GR does not affect the GR localisation but the DNA binding activity of GR, as observed in WT and SIRT2^-/-^ mice. Further, although we have performed immunoprecipitation, immunostaining and dSTORM to show that SIRT2 interacts with GR, we have not conclusively proven direct interaction using rigorous methods such as FRET and protein complementation. Further, in our dex-treated *in vitro* and mouse models, we observed reduced levels of SIRT2, concomitant with increased proteasomal degradation of SIRT2. However, we have not explored the underlying cause of increased SIRT2 degradation under dex-treated conditions. We have also not tested the effect of SIRT2 activation in other models of skeletal muscle degeneration. Nevertheless, our findings indicate that SIRT2 is a critical regulator of muscle homeostasis, and SIRT2 could be a potential therapeutic target for treating muscle atrophy. Future studies are required to develop novel SIRT2 activators for the treatment of muscle atrophy in humans.

### Experimental model and Method details Animal experiments

4-months old C57BL/6J mice were acquired from Central Animal Facility (CAF) IISc. We purchased SIRT2 whole body knock out (SIRT2^-/-^), SIRT2 flox (B6.Cg-Col1a1tm1(CAG-Sirt2)Jmi/DsinJ) and ACTA-Cre mice (FVB.Cg-Tg (ACTA1-care)79Jme/J) mice from Jackson laboratories, USA. Institutional Animal Ethics Committee (IAEC), Indian Institute of Science (CAF/Ethics/833/2021), has approved this project. All the animals were maintained in a well-ventilated, humidity and temperature-controlled environment with 12-hour day and night cycle and kept in IVCs in CAF. Mice were provided with ad libitum food (standard chow diet) and water. Not more than five animals were kept per cage. We generated muscle specific SIRT2-Tg (msSITR2-Tg) by breeding SIRT2 flox with ACTA Cre mice in-house, and the litter was genotyped using gene-specific primers for Transgenic SIRT2 construct and ACTA cre construct. After confirmation, the non-transgenic (N-Tg) and msSIRT2-Tg mice of four months age were used for various experiments. Mice were unbiasedly and randomly divided into various groups within the genotype. Both males and females adult animals were used for the experiments. We performed intraperitonial injection of Dexamethasone at the dose of 10mg/kg/day for 15 days continuously. Their body weight was recorded at intervals of 0, 5,10 and 15 days post-injection. The experiments were terminated after euthanizing the animals using 3 % isofluorane to anesthetize them and performing cervical dislocation. Different muscles (Gastrocnemius, Biceps, Triceps, Quadraceps, Tibialis anterior, Soleus) were weighed. We used vernier callipers to measure tibia bone length for normalization. After collecting the muscles, they were distributed either to snap-freeze in liquid nitrogen or were collected in 10% buffered formalin solution (Sigma Aldrich) for histological experiments. For SUnSET assay [74], on 15th day post Dex injections, the mice were intraperitonially injected with puromycin at a dose of 40nmol/g of body weight. 30 mins post injection, the animals were euthanized, and different muscles were collected and snap-frozen in liquid nitrogen until further use.

### Primary myotube culture

We have followed our previously published protocol to isolate and culture myotubes [75]. Briefly, we have used 0-3 days old neonatal mice or rat pups to isolate primary myotubes. Sex was not considered as a biological variable. The pups were euthanized using 2% isoflurane and then were decapitated. The skin was peeled off from the skeletal muscles (gastrocnemius, quadriceps, and triceps) of the limbs, washed in ice cold PBS with 0.01 M glucose. The isolated muscles were cut into smaller pieces and placed in digestion mixture containing type II collagenase (2 mg per ml of serum-free DMEM). These were placed in a shaker incubator at 250 rpm, 37°C for 90 minutes.After the digestion, the digested myoblasts were collected in serum and pre-plated for 40 minutes to aid the attachment of fibroblasts. The serum containing unattached myoblast cells were subjected to centrifugation at 500G for five minutes. The supernatant was discarded, and the pallet was resuspended in DMEM media (High glucose) containing 10% FBS. The resuspended cells were seeded on gelatin coated culture wares for further experimentation. After 24 hours of incubation, the cells were either supplemented with fresh media or differentiation media based on their confluency. The differentiation media was maintained upto the required time until we observe fully formed myotubes. The plates with mature myotubes were used for experiments. A dose of 50 µM of dexamethasone was used to induce myotube atrophy for 48 hours in serum free media [75]. Following 44 hours of Dex treatment, the primary myotubes were treated with 50 µM of MG132 inhibitor for 4 hours in serum free media and further processed for western blotting. For sea horse experiments, undifferentiated myocytes were used.

### Cell culture

Cells (C2C12 or HEK 293T) were cultured in DMEM high glucose with10% FBS in it along with 1X Antibiotic-Antimycotic solution. Rat glioma cells (C6 cell line, NCCS, Pune) were cultured in Ham’s F12 medium supplemented with 2mM L-glutamine adjusted to contain 90% 1.5 g/L sodium bicarbonate, 9% fetal bovine serum and 1% penicillin–streptomycin at 37 °C. Cells at 80-90% confluency were trypsinized using 0.25% trypsin-EDTA followed by centrifugation at 500 g for 5 minutes, RT. Cells were seeded according to their experimental requirements. Lipofectamine 3000 and p3000 were used for plasmid tranfection whereas RNAimax was used to transfect siRNA as per the manufacturer’s protocol.

### Glucose tolerance test (GTT)

Mice were separated in a fresh cage without food for 12 to 16 hours prior to the start of the GTT experiment. Each time fasting was initiated around 7:00 p.m and the mice were provided with fresh water during fasting. After fasting, the mice were weighed and measured for 0 min blood glucose level was recorded using a glucometer (Accu-Chek Roche) by testing the tail tip blood. Then they were injected with appropriate volume of glucose (2gm/kg of body weight made in 1XPBS) intraperitonially. The blood glucose levels were subsequently measured at the time intervals of 15, 30, 60 and 120 minutes. After terminating the experiment, the animals were provided with food and water and kept back in the colony.

### Insulin tolerance test

To perform Insulin tolerance test (ITT), mice were separated for fasting in a fresh cage without food for 4 to 6 hours before the starting of the experiment. Fasting was initiated at 9:00 a.m., and the mice were provided with fresh autoclaved water. Mice were recorded for their bpdy weight and their 0 min fasting blood glucose levels were recorded using a glucometer (Accu-Chek - Roche) by testing the tail tip blood. The mice were injected with insulin (0.75IU/kg of body weight) and the blood glucose levels were recorded at time intervals of 15,30, 60, and 120 minutes. After termination the experiment, the animals were provided with food and water and placed back in the colony. As a precautionary measure, 20% glucose solution prepared in 1X PBS was kept ready to be injected to rescue the mice suffering from severe hypoglycaemic conditions.

### Blood collection

Mice were anesthetized with 1% isoflurane solution and the blood collection was done with a micro-hematocrit capillary tube through retro-orbital route. The capillary tube tip was pierced near the medial canthus of the right eye below the nictating membrane. By apply gentle force to rotate the tube forward aided in the blood collection from the punctured sinus into the heparin coated tubes. The samples were further processed in veterinary diagnostic lab (Vetlesions and Rohana veterinary diagnostic lab), to test various blood parameters.

### Rotarod performance test

The rotarod performance test was performed based on the previously described protocol [43]. Briefly, the animals were assessed for their ability to balance on a rotating rotarod set to increase its speed gradually. The time any animal takes to fall off of the rotation rod termed as latency, will be recorded and regarded as the measure of muscle function. The instrument contained a clean rod at 30cm height from the instrument base. Prior to recoding the actual data, the animals were trained to walk on a rotarod at a minimal constant speed of 12rpm at 2 sessions for 2 consecutive days. The animals that went under dex injection were trained before injection twice and was recorded for latency measure to avoid internal animal bias and randomly assigned into two groups for vehicle or dex injections. Post injection (after 15 days), the animals were trained once again at training speed and given a day of rest.

Animals were then tested in a single session consisting of 3 trials. Each trial comprised a commencing minimal rotation speed of 12rpm which was gradually subjected to increasing speeds of 20 rpm, 25 rpm, and sustaining at 30 rpm. Each such steps lasted for 20s for a maximum period of 600 seconds. There was a time gap of 15 minutes rest period between each trial. The latency period for each animal was recorded.

### Histology

The tissues samples which were collected in 10% buffered formalin solution after euthanasia are kept in the solution for at least 72 hours for fixation. The formalin was removed by placing the samples in running water for a minimum of 8 hours. The samples are dehydrated in different concentrations of alcohol, then in xylene and embedded in paraffin wax using a tissue processing unit (Leica TP1020, Germany) and made into paraffin blocks. 5µm sample sections were made and collected on glass slides using a microtome (Leica RM2245, Germany). The sections were subjected to H&E staining for muscle fibre morphology, Masson’s trichrome staining for detecting fibrosis and WGA staining (Molecular probes) (5µg/ml) to check myotube cross-sectional area.

### Immunohistochemistry

Gastrocnemius muscle tissue was embedded in OCT solution (Sigme Aldrich) and sectioned 8µm usng cryomicrotome at −15°C (Leica, CM1950, Wetzlar, Germany). For staining the MHC isoforms, the sectioned samples (n=3-4/group) were air dired for 1hour and then rehydrated for 5 mins with 1X PBS. After rehydration, the samples were blocked for 30 mins with the blocking buffer which contains 0.5% BSA and 0.5% triton-X 100 in 1XPBS. The sections were washed 3 times for 1 min each in 5% BSA. Later, the samples were covered and incubated overnight with primary antibody at 40 degree C. Used primary antibodies include Myosin Heavy Chain (MHC) type I antibody (BAD5, DSHB) (1:100), MHC type IIa antibody (SC71, DSHB) (1:100), MHC type Ilb antibody (BFF3, DSHB) (1:100), MHC type IIx antibody (6H1, DSHB) (1:50). Next day, washed the samples 3 times with 5% BSA for 5 mins each. Then, incubated with the secondary antibody corresponding to the respective primary antibody for 1 hour at room temperature. The secondary antibody includes goat anti-mouse IgG (H+L) cross-adsorbed secondary antibody (A11001, Invitrogen), Alexa-FluorTM488 goat anti-mouse IgG (A21121, Invitrogen) (1:500), goat anti-mouse IgM (heavy chain) cross-adsorbed secondary antibody, Alexa-FluorTM595 (A21044, Invitrogen) (1:500). Later, samples were washed with 5% BSA for 3 times for 5 mins each and followed by washing with distilled water once. The slides were mounted using 85% glycerol and cover slips were sealed with transparent nail paint. Images were acquired with Olympus IX 83 inverted microscope (Olympus, IX83, Tokyo, Japan) and the proportion of MHC fibers were calculated.

For Cytochrome oxidase (COX) staining and Succinate dehydrogenase (SDH) staining, the samples were first collected in polyfreeze (Sigma) right after euthanasia and weighing the muscle mass and fixed by placing them on isopentane chilled by using liquid nitrogen and were stored at −80^0^C for further processing [76, 77]. The COX and SDH stains were made used to differentiate among different fiber types in the muscle tissue. The sections were cut at 8µm sample sections using a cryomicrotome (Leica, CM1950, Germany) maintained between −15 to −20°C, on glass slides and were air dried for 30 minutes. For COX mitochondrial staining, the slides were soaked in solution containing 100µM cytochromeC in 0.1M PBS (of pH 7) (Sigma), and 2µg/ml of bovine catalase (Sigma), 1X DAB (Sigma). The slides were then washed 3 times in 0.1MPBS for 10 minutes per wash. For SDH staining, the slides were soaked in staining-solution (SRL, India) having 0.1M Magnesium chloride, 2.4 mM tetrazolium and 0.2M succinic acid in 0.2M PBS (pH 7.4). After incubation the slides were washed for 3 minutes in DI water twice. The slides were then dried and mounted with Distyrene Plasticizer and Xylene (DPX) and proceeded for further imaging.

### Quantitative RT-PCR

RNA isolation was performed using phenol-chloroform extraction method (RNAiso plus from Takara). A maximum of 0.5µg of RNA was used for cDNA synthesis using the 5X iScript cDNA synthesis kit (Bio-Rad). Prepared cDNA was used to check the expression of different genes by qRT-PCR using iTaq Universal SYBR Green Supermix (Bio-Rad). QuantStudio 6 Flex System (Thermo Fisher Scientific) was used for qRT-PCR reaction. The primer sequences are listed in Table S1.

### RNA-seq data analysis

Adapters in the raw sequencing reads were trimmed using fastp 0.23.4 (modified parameters: --detect_adapter_for_pe --qualified_quality_phred 20 –correction), aligned to the GRCm39 genome using star 2.7.10b (default parameters) and using samtools 1.16.1, the resulting .sam files were converted .bam files and filtered to retain only paired-end reads (modified parameter: -bf 1). Raw read counts were generated using subread 2.0.1 featureCounts (default parameters). Differential gene expression analysis was performed in R with edgeR 4.2.2, and volcano plots were generated using ggplot21 3.5.1. Genes with FDR-adjusted p-values < 0.05 were called as differentially expressed. GO analysis of the upregulated and downregulated differentially expressed genes was carried out using Metascape2, enriched GOs were plotted with ggplot2 and heatmaps of the z-scored counter per million scaled raw read counts of the individual genes in each replicate belonging to enriched GOs were generated using ComplexHeatmap3 2.20.0. [78–80]. The transcriptome of WT and SIRT2^-/-^ gastrocnemius skeletal muscles samples have been deposited at NCBI SRA (BioSample accessions: SAMN47809844, SAMN47809845, SAMN47809846, SAMN47809847, SAMN47809848, SAMN47809849)

### Histogram and UMAP of SIRT2 and GR

The data were analyzed from the Muscle Aging Cell Atlas [45], which contains single-cell RNA-seq and single-nucleus RNA-seq data from patients. Briefly, the data consists of intercostal muscles isolated from 17 individuals. We then selected SIRT2 and GR expression in the single-cell and single-nucleus data and saw their expression profile compared against one another.

### Oxygen consumption and lactate production (Seahorse mitostress test)

To determine the differences in oxygen consumption rate (OCR) and extracellular acidification rate (ECAR) under SIRT2-depleted conditions, we cultured untreated control, and AGK2-treated (24h treatment) primary rat myocytes, as well as wild-type and SIRT2^-/-^ mouse muscle-derived primary myocytes independently. These primary myocytes were subjected to Seahorse XF Mito Stress Test according to the manufacturer’s instructions (Agilent Technologies). 30,000 cells were seeded per well in an 8-well Seahorse flux analyzer plate precoated with Cell-Tak (Corning). Cells were incubated for 1 h at 37°C in a non-CO_2_ incubator before starting the assay. Basal OCR and ECAR were determined with readings at three time points under unstimulated conditions, followed by treatment with 1 µM oligomycin, 1 µM carbonyl cyanide-4-(trifluoromethoxy) phenylhydrazone (FCCP), and 0.75 µM rotenone + antimycin A. Wave 2.6.3 software (Agilent Technologies) was used for mitochondrial respiration parameter analysis, and data were normalized to total protein content per well.

### Western blot

Frozen muscles tissues were crushed into powder by motor and pestle to make protein lysates using RIPA lysis buffer (50 mM Tris-HCl pH7.8, 150mM NaCl, 1mM EGTA, 1mM EDTA, 0.5% SDC, 1% SDS, sodium orthovanadate, NaPP, Phenylmethylsulfonyl fluoride, and protease inhibitor cocktail). Same buffer composition was used to prepare cell lysates as well. After vertaxing, the samples were subjected to centrifugation at 15000 rpm, 4°C for 15 minutes. After collecting the lysates, protein concentration assessed using Bradford (Bio-Rad) assay. After normalizing, equal amount of protein was loaded on the SDS-PAGE at 80 volts for 2 hours. The gel was set for overnight wet transfer on a PVDF membrane at 20 volts, 4°C. The membrane was blocked for 1 hour with 5% skimmed milk made in Tris-buffered saline in 0.1% Tween 20 and later used to probe with primary antibody prepared in 5% BSA for 16 hours at 4°C. Specifically bound primary antibody was detected by incubating the membrane with secondary antibody conjugated with horseradish peroxidase (HRP) and later developed using ECL reagent (Bio-Rad) in a chemidoc (Bio-Rad).

### Co-immunoprecipitation assay

HEK 293T or C1C12 cells were washed with 1XPBS for 2-3 times and scraped with RIPA buffer (150mM NaCl, 50mM Tris-HCl, 1% NP-40, 0.1%SDS, 0.25% sodium deoxycholate, 1mM EDTA, 2.5mM sodium pyrophosphate, 1mM Na3VO4, protease inhibitor cocktail, 1mM PMSF, 1µM ADP-HDP) maintained at 4°C. The cells mixed with lysis buffer were vortexed vigorously for 15 seconds, followed by 5 minutes at 4°C for 30 minutes. The crude lysate was further centrifuged at 12,000 G for 10 minutes at 4°C. The supernatants were collected in new micro centrifuge tubes, and the protein concentration was measured using Bradford assay. 0.5mg of total protein was incubated with 2µg of various antibodies for overnight incubation at 5rpm at 4°C. After 16 hours incubation, the protein G Magnetic beads (Med Chem express) were washed three times with RIPA buffer and incubated with specific protein antibody complex at 5rpm at 4°C for 2 hours. After the incubation, beads were washed thrice with RIPA buffer and then incubated with 2X laemmli buffer (Bio-Rad) for 5 minutes at 95°C to elute the protein in laemmli buffer. The elution was then validated for interaction after immunoblotting using various antibodies.

### Chromatin immunoprecipitation (ChIP) assay

ChIP assay was performed using mice gastrocnemius muscles of wild type and SIRT2 knock out animals using the commercially available kit as per the manufacturer’s protocol (Cell Signaling). Briefly, the tissues were first homogenized and cross-linked with 1% formaldehyde solution for 15 minutes at RT and then quenched using 0.15 M glycine for 10 minutes. The tissue suspension was further homogenized by Dounce homogenizer and the pellet containing the chromatin was sheared using micrococcal nuclease, followed by cycles of sonication. The sheared chromatin in the range of 100 to 800bp was use for immunoprecipitating crosslinked DNA-protein complex. GR antibody was used for IP. H3 antibody and normal rabbit IgG antibody were used as positive and negative control respectively. The eluted DNA-protein complex and 10% input chromatin for each sample were then decrosslinked for 2 hours following which DNA isolation and qPCR were performed using specific primers.

### Fluorescence microscopy

Upon the completion of treatment/transfection, cells were fixed with 3.7% formaldehyde at RT for 15 minutes, followed by permeabilization with 0.2% triton X100 and then blocking with 5% BSA in PBST (1XPBS with 0.1% Tween 20). Further, cells were washed with PBST for 3 minutes at RT three times and incubated with specific primary antibodies at 4°C for 16 hours. After primary antibodies incubation, cells were further incubated with fluorophore-conjugated secondary antibodies (Alexa Fluor 594 and 488). Hoechst was used to stain nucleus and then mounted using fluoromount reagent (Sigma). Images were acquired using confocal microscope (Carl-Zeiss LSM 710/880, Andor Dragonfly 302 Spinning Disk Confocal) [51, 81].

### Direct Stochastic Optical Reconstruction Microscopy (dSTORM) imaging and analysis

For dSTORM imaging, C6 rat glioma cells after 24 hours of transfections, were fixed using 4% paraformaldehyde and 4% sucrose in PBS for 10 min at 4 °C, quenched with 0.1 M glycine in PBS at room temperature, permeabilized with 0.25% TritonX-100 for 5 min and blocked with 10% BSA in PBS solution for 30 min at room temperature. The cells were incubated with the primary antibodies (Anti FLAG-Rabbit (SIRT2), Anti GFP-Mouse (GR)) diluted in 3% BSA (1:200) for 1 h at room temperature in a humidified chamber. The cells were washed four times (5 min each) with 3% BSA and then incubated with suitable secondary antibodies (anti-rabbit 642 and anti-mouse 532) for 45 min followed by 4-time washes of 5 min each. The 12-well plates were wrapped in an aluminum foil and stored in PBS at 4 °C for dSTORM imaging. Imaging was done within 24 h of completion of labeling procedure.

dSTORM imaging was conducted at 37°C in an open chamber (Ludin chamber, Life Imaging Services) mounted on an inverted motorized microscope (Olympus IX83) equipped with a 100× TIRF objective, NA 1.49 allowing extended acquisition by TIRF illumination (Total Internal Reflection Fluorescence). MetaMorph software was used to control illumination and acquisition. 100 nm diameter beads were incubated on coverslips for 30 minutes (Tetraspeck; Thermo Fisher Scientific, USA) and used as fiduciary marker for lateral drift correction. Immunostained cells were imaged in a dSTORM buffer containing catalase, TCEP, glycerol, glucose, and glucose oxidase dissolved in a Tris-HCl buffer, used to induce random molecule activation [82]. Photoconversion of Alexa-647 and Alexa-532 fluorophores were performed by illuminating the sample with respective excitation lasers at 300 mW. Five sets of 4000 images were obtained with an exposure time of 20 ms, acquiring a total of 20,000 images [83]. Single molecule fluorescence was collected using a sensitive EMCCD camera (Evolve, Photometrics). Acquisition was controlled by MetaMorph software (Molecular Devices).

#### Image analysis

SIRT2 and GR protein single molecules present in the cell were identified and quantified from super-resolution images using the Palm Tracer plugin in Metamorph software [84]. Chromatic aberration correction and lateral drift correction was done on PalmTracer using 100nm sized fiducial markers to ensure the single molecule level co-localization of the two proteins at nanoscale. Images from the 642(SIRT2) and 532(GR) channels were then color aligned on metamorph. Colocalization of SIRT2 and GR was visible at this point. To further quantify the co-localization and look at the difference in interaction between WT and NA groups, ROIs of size 250*250 pixels were created on each image and colocalization analysis was done on these cropped regions on ImageJ using coloc2 plugin. Similar paradigms were used to quantify the co-localization between SIRT2 and GR in Wild Type vs PCDNA control in confocal images.

### Luciferase assay

We performed Glucocorticoid receptor activity assay using GRE-Luc (MMTV-Luv) plasmid as described previously [85]. Briefly, we co-transfected GRE-Luc with renilla plasmid (pRLCMV) for hours. Lysates were prepared using passive lysis buffer (using Promega’s dual luciferase kit) as per manufacturer’s protocol, and luciferase activity was assessed using a luminometer.

### Protein-Protein Docking

The protein-protein docking was performed using the HADDOCK software. The crystal structures of SIRT2 monomer (PDB code: 5Y0Z) and the ligand binding domain of GR (PDB code: 7PRX) were used as input for the docking process. HADDOCK utilizes a hybrid approach combining information-driven docking and molecular dynamics simulations to generate the docked complex structures. After the docking simulations, the resulting structures were analyzed using HADDOCK’s scoring functions and clustering algorithms. The top cluster, representing the most favourable and consistent docking solutions, was selected for further analysis. The selected docked complex structure was visualized and analyzed using UCSF Chimera, a molecular visualization software. The hydrogen bonds formed between the residues of SIRT2 and GR in the selected docked complex were identified and analyzed using UCSF Chimera. The presence of hydrogen bonds was determined based on distance and geometry criteria.

### *In-silico* acetylation experiment

The Protein Data Bank (PBD) currently does not have a record of the full-length structure of the human glucocorticoid receptor (GR), consisting of an N-terminal domain, a DNA-binding domain, and a ligand-binding domain. However, 17 crystal structures of the DNA-binding domain of GR complexed to DNA fragments are available. Of these, PDB code 5E6C PDB [86] was used for the in-silico acetylation-based study. This structure has a resolution of 2.20 Å, includes all lysine residues of interest, and lacks any mutations. Further, it is in the dimer, which is the biologically relevant form of GR. In-silico acetylation was performed using the PyMOL plugin, PyTMs [87]. A total of 9 lysine residues, 5 in chain A and 4 in chain B, were acetylated. Each acetylated lysine residue is represented as ALY.

### Energy minimization

Both unacetylated and acetylated human GR DNA-binding domains complexed to genomic DNA were subjected to energy minimization using CHARMM27 force field and the steepest descent minimization algorithm available in GROMACS [88]. Following this, protein-DNA interaction network analysis was performed. A donor-acceptor distance of <3.5 Å was set as a starting point for detecting potential hydrogen bonds. The hydrogen bonds formed between the DNA and the protein were then compared between the acetylated and unacetylated forms of human GR using the UCSF Chimera visualization module [89].

### Normal Mode Analysis

Using a Bio3D R package, atomic fluctuations from an all-atom Normal Mode Analysis (NMA) were calculated to study the flexibility associated with the protein-DNA complex before and after GR acetylation [90]. The top 10 slowest modes, which provide information on the global motions of a protein, were used to demonstrate the differences in flexibility between the unacetylated and acetylated forms of human GR complexed to DNA. The difference in fluctuations for all residues of GR in both forms was calculated and plotted.

### Quantitative image and data analysis

After acquiring the fluorescence images, ZEN black (Carl-Zeiss), Metamorph and ImarisViewer 10.0.1 softwares were used to process them. All the image quantifications were done using ImageJ Fiji software (National Institutes of Health, USA). The western blot images were processed using Image Lab (Bio rad) software. Adobe Photoshop CS software was used to crop the images. For qRT-PCR analysis, QuantStudio Real-Time PCR software was employed. The images or graphs were then arranged in Adobe illustrator CS4. Metamorph software, Wave 2.6.3 software (Agilent Technologies), was used to analyse dSTORM data.

### Statistical analysis

All the graphs were prepared and analyzed for data normality and appropriate statistical tests in Graphpad Prism 8.4.2 software. All the analyzed data are represented as mean ± SD. Kolmogorov-Smirnov and Shapiro-Wilk tests were used to determine data normality. Appropriate statistical analysis was adapted to analyze the data: all normally distributed data were analyzed by applying Student’s unpaired t-test or one-way analysis of variance (ANOVA) with Tukey’s multiple comparisons test and two-way ANOVA with Tukey’s multiple comparisons test. Mann–Whitney test or Kruskal–Wallis test with Dunn’s multiple comparisons were used for non-normally distributed data set. Differences in mean values between groups were considered as statistically significant at a two tailed p-value ≤ 0.05.

### Study Approval

Institutional Animal Ethics Committee (IAEC), Indian Institute of Science (CAF/Ethics/833/2021), has approved this project.

## ACKNOWLEDGEMENTS

The Central Animal Facility, Central Confocal facility (Division of Biological Sciences and Department of Microbiology and Cell Biology), and the microtome facility (Division of Biological Sciences), Advanced User Facility (Centre for Neuroscience), all at the Indian Institute of Science, Bengaluru, and the cryo-microtome facility and the fluorescent microscope facility (Department of Biotechnology) at the Indian Institute of Technology Madras, Chennai are acknowledged for their services and the technical help.

## Author Contribution

NRS conceived and designed the study and prepared the final draft. AKT designed the experiments. AKT and BS analysed the experiments. AKT, BS, AM, AB, SR and AJS wrote the manuscript. SN, AKT, BS and DR performed the primary myotubes culture and confocal microscopy. AKT, BS, AB, DN and AJS did myotubes quantification. BS and AKT performed the nuclear localization experiments. AKT and BS bred the SIRT2 knockout and developed the msSIRT2-transgenic mice line. BS and AKT performed genotyping of the SIRT2 knockout and msSIRT2-transgenic mice line. AKT, MS and BS performed glucose tolerance tests and insulin tolerance tests. AKT, SN and BS performed the qPCR. ABP, DR, AB, DN, MBM, BS and AKT performed immunoblotting. RS designed and conceived computational structural analyses. SS, HT wrote the results for computational analysis experiments. AKT, PAD and BS performed histological experiments. ASN performed RNA-seq data analysis. AKT, PAD, and SN acquired images for the histological section. AKT and BS quantified the confocal images. DS and NAJ performed the COX, SDH and MHC fiber typing experiments. AKT and BS performed Co-Immunoprecipitation experiments. ABP and VSR performed mass spectrometry-based experiments and site-directed mutagenesis of the Glucocorticoid receptor. AKT, BS, AB and LD conducted rotarod experiments. AJS and DN performed the luciferase reporter assay. ABP, DN, AJS and BS performed SUnSET assay. BS and AB conducted MG132 experiment. BS, SN and SR performed ChIP experiments. MKER and DN performed dSTORM experiment. HMJ, ABP, AM and AS performed seahorse experiments. RM is involved in the project discussion. Original data have been cross verified by SR, BS, AKT, AJS and NRS. AKT and BS are co first authors. Initial experiments were carried out by AKT, the project was later managed by BS.

## Data and code availability

RNA sequencing data that support the findings of this study, have been deposited in Sequence Read Archive with the sample accession codes: SAMN47809844, SAMN47809845, SAMN47809846, SAMN47809847, SAMN47809848, SAMN47809849. Source data supporting the findings in this study underlying dot plots, violin plots, and uncropped western blots are provided in the excel raw data file.

## Materials availability

All unique/stable reagents generated in this study are available from the Lead Contact with a completed Materials Transfer Agreement.

## Lead Contact

Further information and requests for resources and reagents should be directed to and will be fulfilled by the Lead Contact, N. Ravi Sundaresan.

## Supplementary Figures

**Supplementary figure 1.**
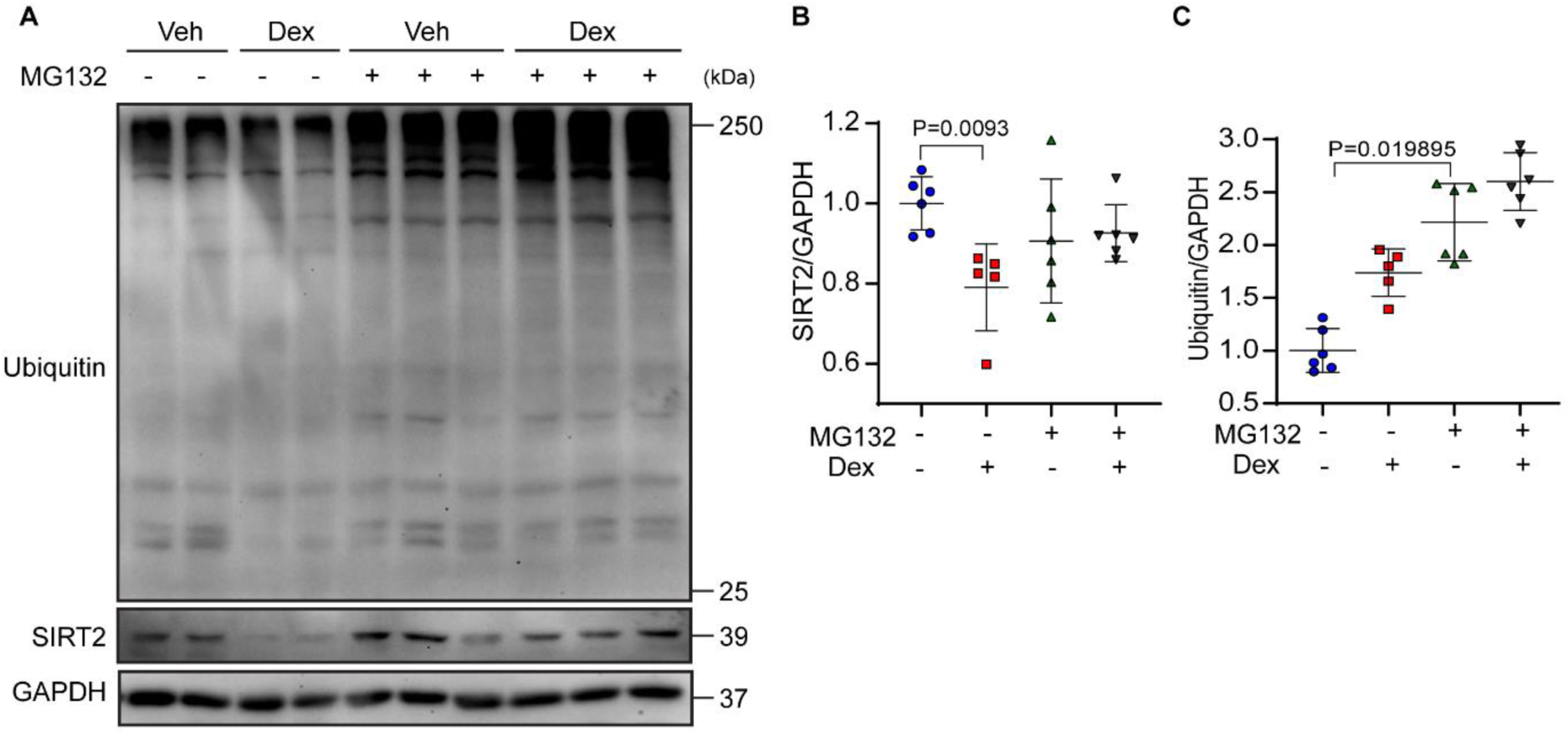
SIRT2 expression in Dex-treated myotubes **(A)** Representative immunoblot showing SIRT2 and ubiquitin levels in the dex-treated primary myotubes under MG132 treatment. Indicated sizes are in kilodaltons (kDa). *n* = 5-6. **(B)** Quantification of SIRT2 levels using immunoblot shown in Supplementary Figure 1(A). Protein levels were normalized with GAPDH. Data represented as mean ± s.d., *n* = 5-6. **(C)** Quantification of Ubiquitin levels using immunoblot shown in Supplementary Figure 1(A). Protein levels were normalized with GAPDH. Data represented as mean ± s.d., *n* = 5-6. Kruskal-Wallis test with Dunn’s multiple comparisons (B, C) statistical tests were used.

**Supplementary figure 2.**
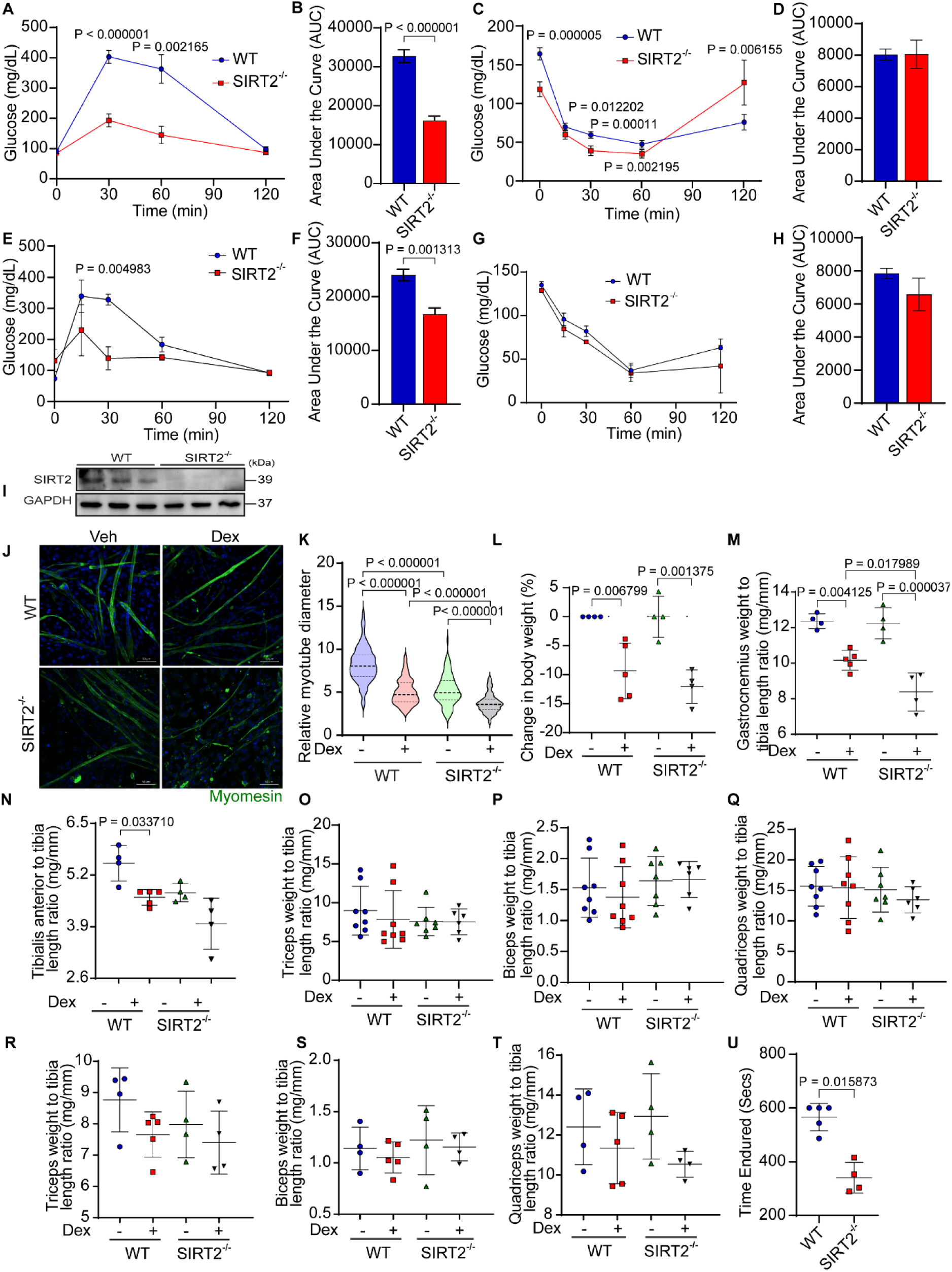
Assessment of Glucose and insulin sensitivity in SIRT2^-/-^ mice **(A)** Quantification of glucose levels showing glucose tolerance test (GTT) in wild-type (males) and SIRT2^-/-^ mice(males). Data represented as mean ± s.d., n = 6. **(B)** Quantification of the area under the curve for Figure S2A in wild-type and SIRT2^-/-^ mice (males). Data represented as mean ± s.d. **(C)** Quantification of glucose levels showing insulin tolerance test (ITT) in wild-type and SIRT2^-/-^ mice (males). Data represented as mean ± s.d., n = 6. The student’s t-test with Welch’s correction was used for statistical analysis. **(D)** Quantification of the area under the curve for Figure S2C in wild-type and SIRT2^-/-^ mice(males). Data represented as mean ± s.d. **(E)** Quantification of glucose levels showing glucose tolerance test (GTT) in wild-type (females) and SIRT2^-/-^ mice (females). Data represented as mean ± s.d., n = 4-5. **(F)** Quantification of the area under the curve for Figure S2E in wild-type and SIRT2^-/-^ mice (females). Data represented as mean ± s.d. **(G)** Quantification of glucose levels showing insulin tolerance test (ITT) in wild-type and SIRT2^-/-^ mice (females). Data represented as mean ± s.d., n = 4-5. The student’s t-test with Welch’s correction was used for statistical analysis. **(H)** Quantification of the area under the curve for Figure S2G in wild-type and SIRT2^-/-^ mice (females). Data represented as mean ± s.d. **(I)** Representative immunoblot to verify SIRT2 deficiency in primary myotubes isolated from SIRT2^-/-^ mice. Indicated sizes are in kilodaltons (kDa). **(J)** Representative immunofluorescence images showing myotubes diameter in primary myotubes isolated from wild-type and SIRT2^-/-^mice when treated with vehicle or dex. Myotubes were stained with antibodies against myomesin (green). Scale bar = 50 µm. **(K)** Quantification of relative myotube diameters in dex-treated primary mice myotubes under SIRT2 deficient conditions, normalized with the vehicle-treated primary myotubes of wild-type mice. Data represented as median with 25 and 75 percentile *n* = 3, N= 58-122. **(L)** Change in percentage body weight of SIRT2^-/-^ mice (female) compared to wild-type mice (females). after 15 days of dex injection (10mg/kg/day) compared to the vehicle group. Data represented as mean ± s.d., n = 4-5. **(M)-(N)** Gastrocnemius, tibialis anterior weight to tibia length ratio in vehicle and dex(10mg/kg/day) injected wild-type mice (females) and SIRT2^-/-^ mice (females). Data represented as mean ± s.d. n = 4-5. **(O)-(Q)** Biceps, triceps, and quadriceps weight to tibia length ratio in vehicle and dex(10mg/kg/day) injected wild-type and SIRT2^-/-^ mice (males). Data represented as mean ± s.d. n = 7-8. **(R-T)** Biceps, triceps, and quadriceps weight to tibia length ratio in vehicle and dex(10mg/kg/day) injected wild-type and SIRT2^-/-^ mice (females). Data represented as mean ± s.d. n =4-5. **(U)** Quantification of time the wild-type and SIRT2^-/-^ mice (female) endured in the rotating rod. Data are shown as mean ± s.d., n =4-5. The two-tailed unpaired student’s t-test with Welch’s correction for test gtt 30min significance and Two-tailed Mann-Whitney statistical test for gtt 60 min significance (A) The two-tailed unpaired student’s t-test with Welch’s correction test for itt 15, 30, and 60min significance (C, E, G) Kruskal-Wallis test with Dunn’s multiple comparisons (K, L, O) Two-way ANOVA with Tukey’s multiple comparisons test (M, N, P, Q, R, S, T) Two-tailed Mann-Whitney statistical test (U) was used for statistical analysis.

**Supplementary figure. 3.**
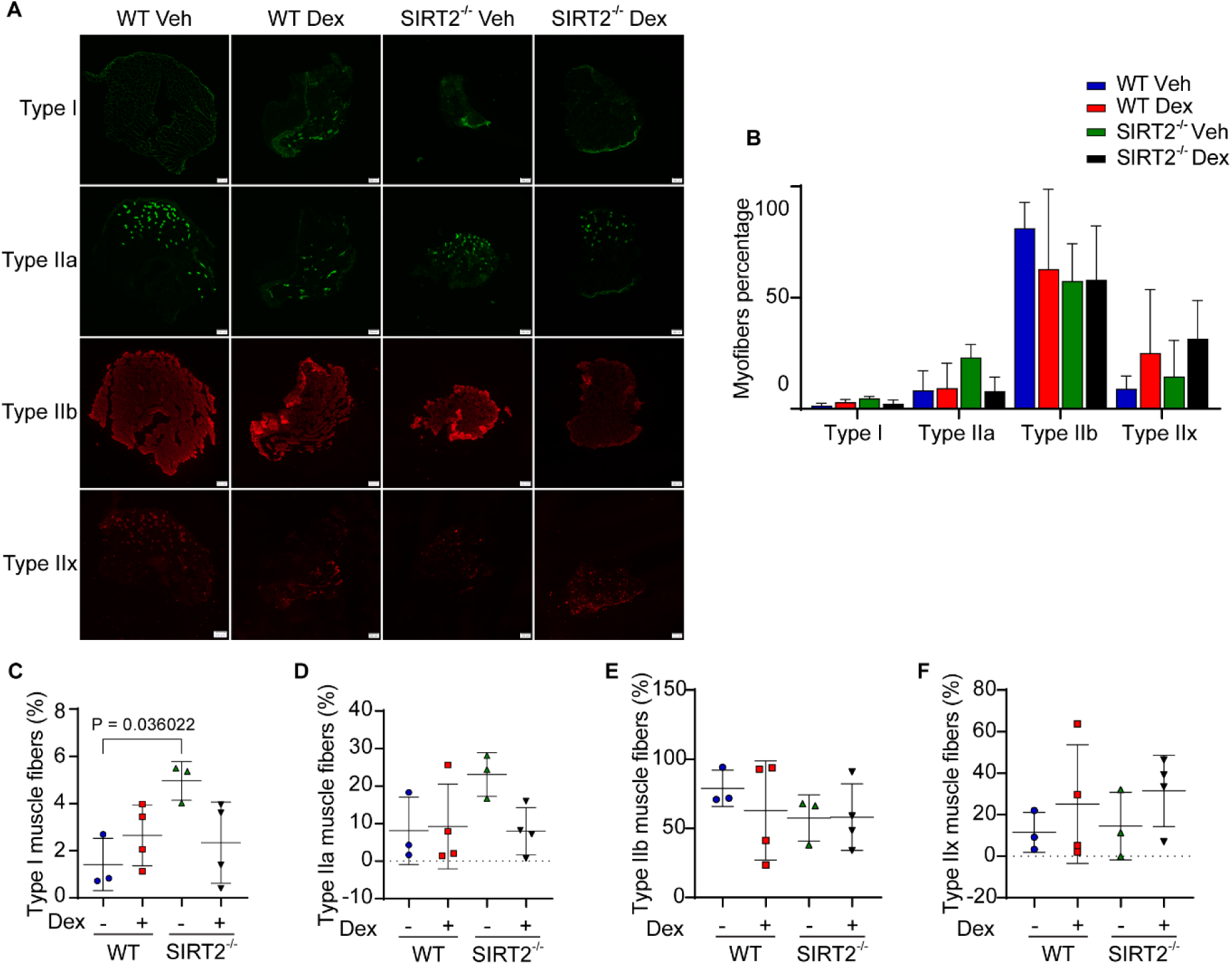
Muscle fibre type distribution in SIRT2^-/-^ mice **(A)** Representative immunofluorescence images showing various muscle fiber types (Type I, IIa (green) and Type IIb, IIx (red). Antibodies for Type I, IIa, IIb, IIx were probed in the gastrocnemius sections of wild-type and SIRT2^-/-^ mice administered with either vehicle or dex(10mg/kg/day). Scale bar = 200 µm. **(B)** Bar graph showing the percentage of various fiber types in the gastrocnemius muscles of SIRT2^-/-^ mice injected with dex. Data represented as mean ± s.d., n = 3-4. **(C)-(F)** Scatterplot graphs representing the percentage of type I, IIa, IIb, and IIx as shown in Figure S3A in the gastrocnemius muscles of SIRT2^-/-^ mice administered with dex. Data represented as mean ± s.d., n = 3-4. Two-way ANOVA with Tukey’s multiple comparisons was used for statistical analysis (C, D, E, F)

**Supplementary figure 4.**
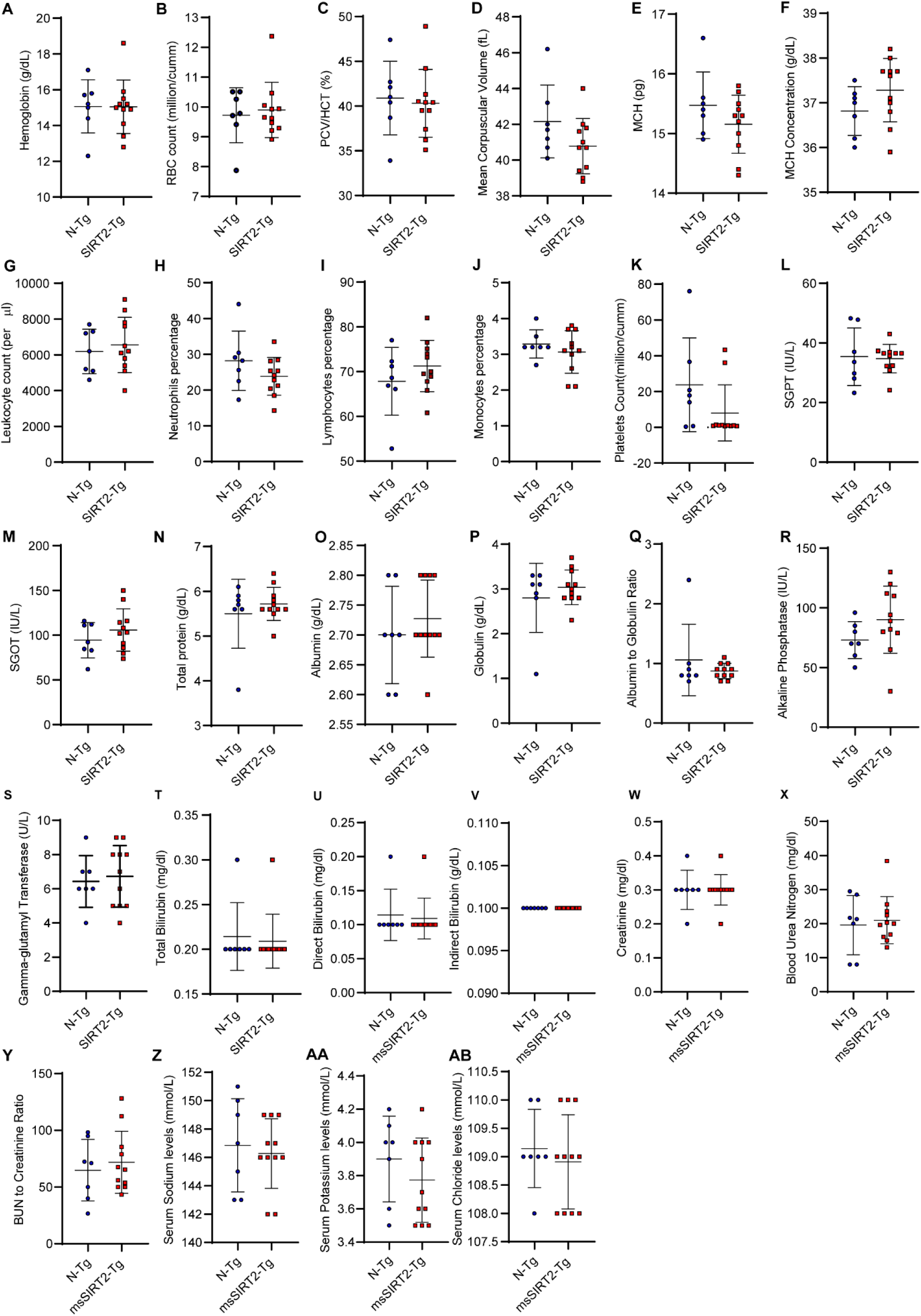
Blood parameters in the msSIRT2-transgenic mice (male) **(A)-(AB)** Quantification of hemoglobin, Red Blood Corpuscles (RBC) count levels, hematocrit content (PCV/HCT), RBC size(Mean Corpuscular Volume), hemoglobin content (MCH), hemoglobin concentration (MCHC), leukocyte count, percentage of neutrophils, percentage of lymphocytes, monocytes, platelets count, alanine aminotransferase concentration (SGPT), aspartate aminotransferase concentration (SGOT), total protein content, albumin concentration, globulin concentration, albumin to globulin ratio, alkaline phosphatase concentration, gamma-glutaryl transferase concentration, total bilirubin concentration, direct bilirubin concentration, indirect bilirubin concentration levels, creatinine levels, Blood Urea Nitrogen(BUN) concentration levels, BUN to creatinine ratio, sodium, potassium, and chloride levels in non-transgenic control and msSIRT2 transgenic (Male). Data represented as mean ± s.d., *n* = 7-11. The two-tailed unpaired student’s t-test with Welch’s correction (A,C,D,E,F,G,H,I,J,K,L,M,N,O,P,Q,R,S,X,Y,AA) Two-tailed Mann-Whitney statistical test (B,T,U,V,W,Z,AB,AC) was used for analysis.

**Supplementary figure 5.**
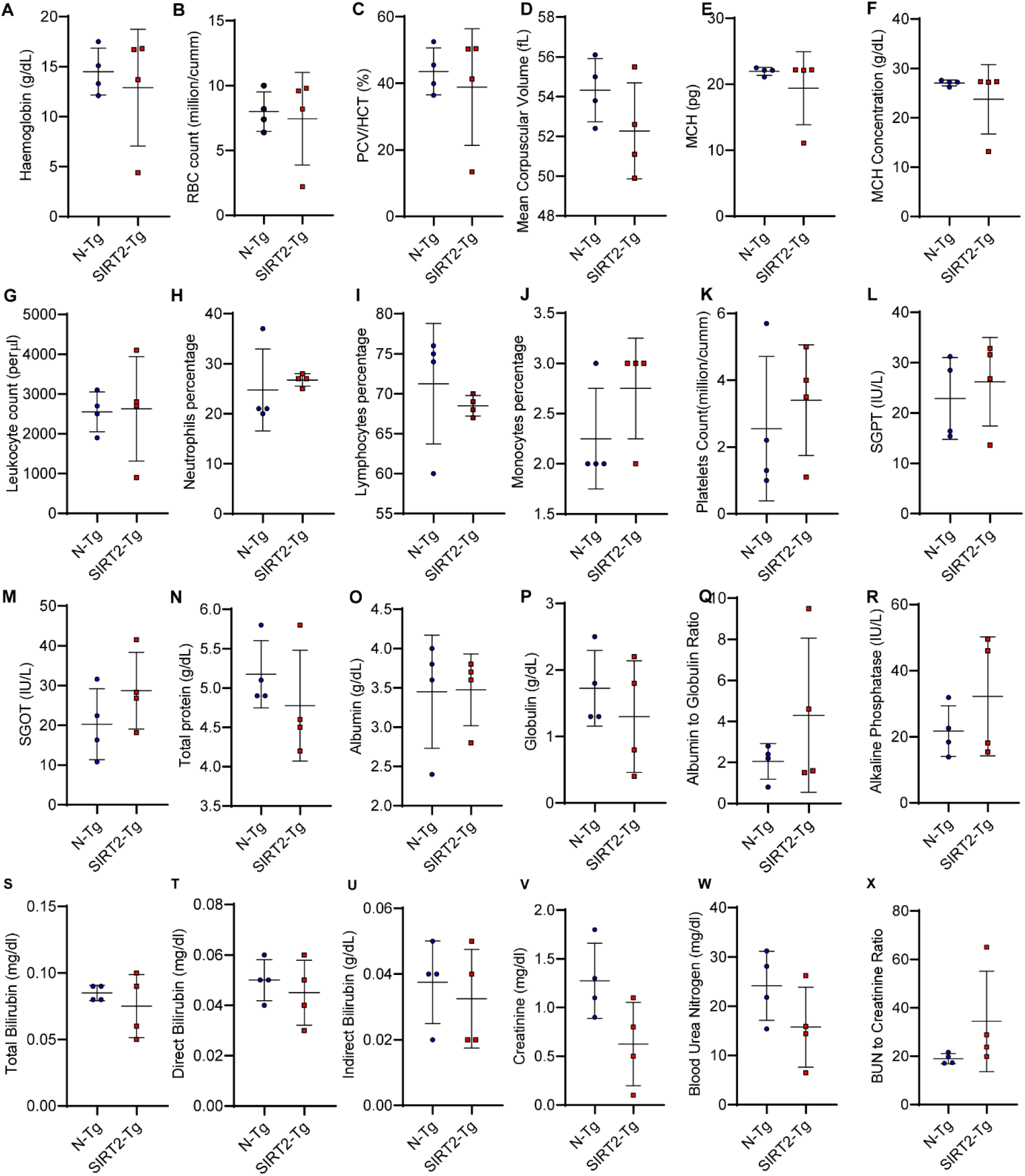
Blood parameters in the msSIRT2-transgenic mice (female) **(A)-(X)** Quantification of hemoglobin, Red Blood Corpuscles (RBC) count levels, hematocrit content (PCV/HCT), RBC size(Mean Corpuscular Volume), hemoglobin content (MCH), hemoglobin concentration (MCHC), leukocyte count, percentage of neutrophils, percentage of lymphocytes, monocytes, platelets count, alanine aminotransferase concentration (SGPT), aspartate aminotransferase concentration (SGOT), total protein content, albumin concentration, globulin concentration, albumin to globulin ratio, alkaline phosphatase concentration, total bilirubin concentration, direct bilirubin concentration, indirect bilirubin concentration levels, creatinine levels, Blood Urea Nitrogen(BUN) concentration levels, BUN to creatinine ratio in non-transgenic control and msSIRT2 transgenic (Female). Data represented as mean ± s.d., *n* = 4. The two-tailed unpaired student’s t-test with Welch’s correction (A,C,D,G,K,L,M,N,O,P,Q,R,S,T,U,V,W,X) Two-tailed Mann-Whitney statistical test (B,E,F,H,I,J)

**Supplementary figure 6.**
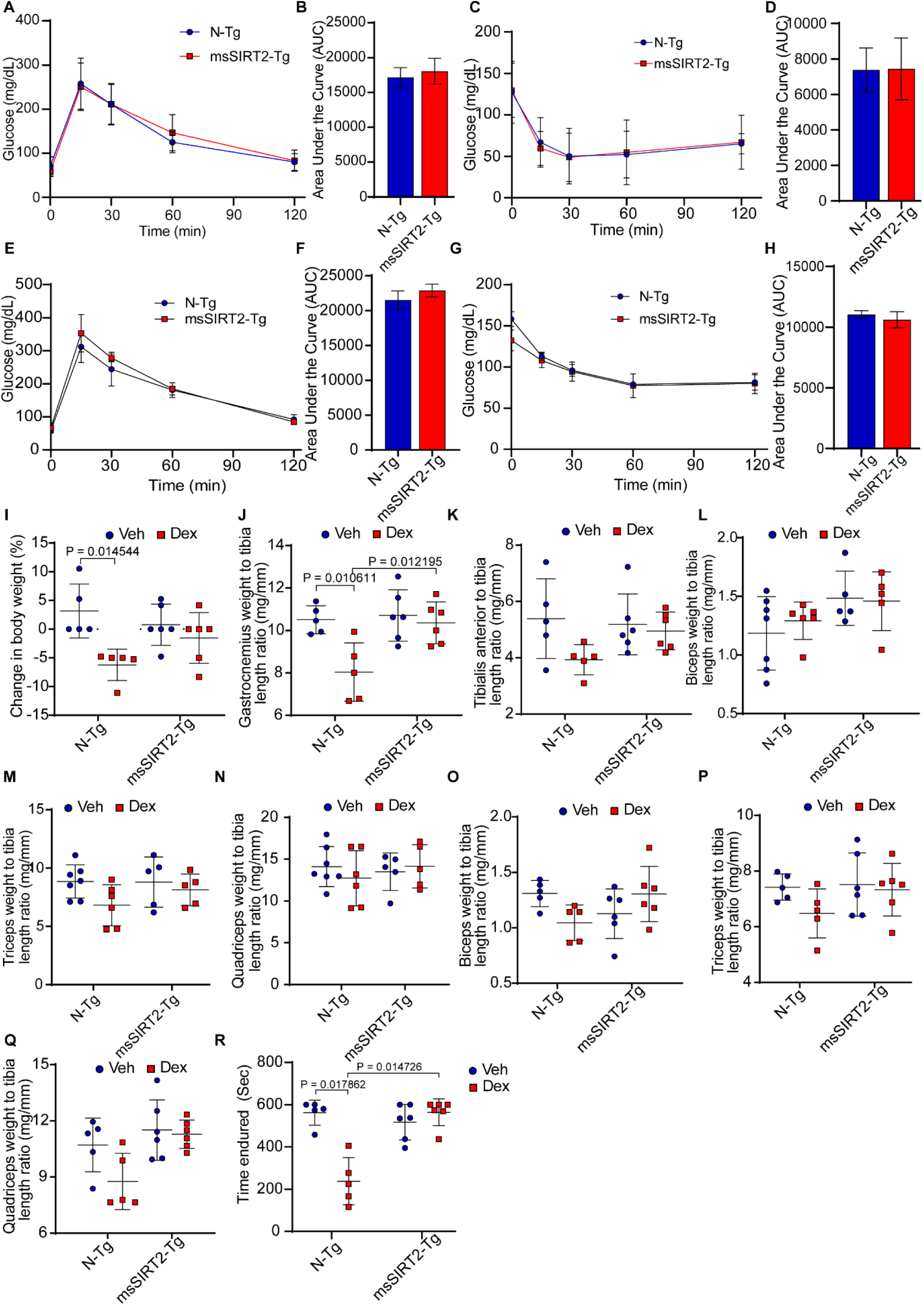
Assessment of Glucose and insulin sensitivity in the msSIRT2 transgenic mice **(A)** Quantification of glucose levels showing glucose tolerance test (GTT) in non-transgenic (males) and msSIRT2 transgenic mice (males). Data represented as mean ± s.d., n = 6. **(B)** Quantification of the area under the curve for Figure S6A in non-transgenic and msSIRT2 transgenic mice (males). Data represented as mean ± s.d. **(C)** Quantification of glucose levels showing insulin tolerance test (ITT) in non-transgenic and msSIRT2 transgenic mice (males). Data represented as mean ± s.d., n = 6. The student’s t-test with Welch’s correction was used for statistical analysis. **(D)** Quantification of the area under the curve for Figure S6C in non-transgenic and msSIRT2 transgenic mice (males). Data represented as mean ± s.d. **(E)** Quantification of glucose levels showing glucose tolerance test (GTT) in non-transgenic (females) and msSIRT2 transgenic mice (females). Data represented as mean ± s.d., n = 3. **(F)** Quantification of the area under the curve for Figure S6E in non-transgenic and msSIRT2 transgenic mice (females). Data represented as mean ± s.d. **(G)** Quantification of glucose levels showing insulin tolerance test (ITT) in non-transgenic and msSIRT2 transgenic mice (females). Data represented as mean ± s.d., n = 3. The student’s t-test with Welch’s correction was used for statistical analysis. **(H)** Quantification of the area under the curve for Figure S6G in non-transgenic and msSIRT2 transgenic mice (females). Data represented as mean ± s.d. **(I)** Change in percentage body weight of msSIRT2 transgenic (female) compared to non-transgenic mice (females). after 15 days of dex injection (10mg/kg/day) compared to the vehicle group. Data represented as mean ± s.d., n = 5-6. **(J)-(K)** Gastrocnemius and tibialis anterior weight to tibia length ratio in vehicle and dex(10mg/kg/day) injected non-transgenic (females) and msSIRT2 transgenic mice (females). Data represented as mean ± s.d. n = 5-6. **(L)**-**(N)** Biceps, triceps, and quadriceps weight to tibia length ratio in vehicle and dex(10mg/kg/day) injected non-transgenic and msSIRT2 transgenic mice (males). Data represented as mean ± s.d. n = 5-7. **(O)-(Q)** Biceps, triceps, and quadriceps weight to tibia length ratio in vehicle and dex(10mg/kg/day) injected non-transgenic and msSIRT2 transgenic mice (females). Data represented as mean ± s.d. n = 5-6. **(R)** Quantification of time the non-transgenic and msSIRT2 transgenic mice (female) endured in the rotating rod. Data are shown as mean ± s.d., n =5-6. Two-way ANOVA with Tukey’s multiple comparisons (J, K, M, N, O, P, Q) Kruskal-Wallis test with Dunn’s multiple comparisons (I, L, R) was used for statistical analysis.

**Supplementary figure 7.**
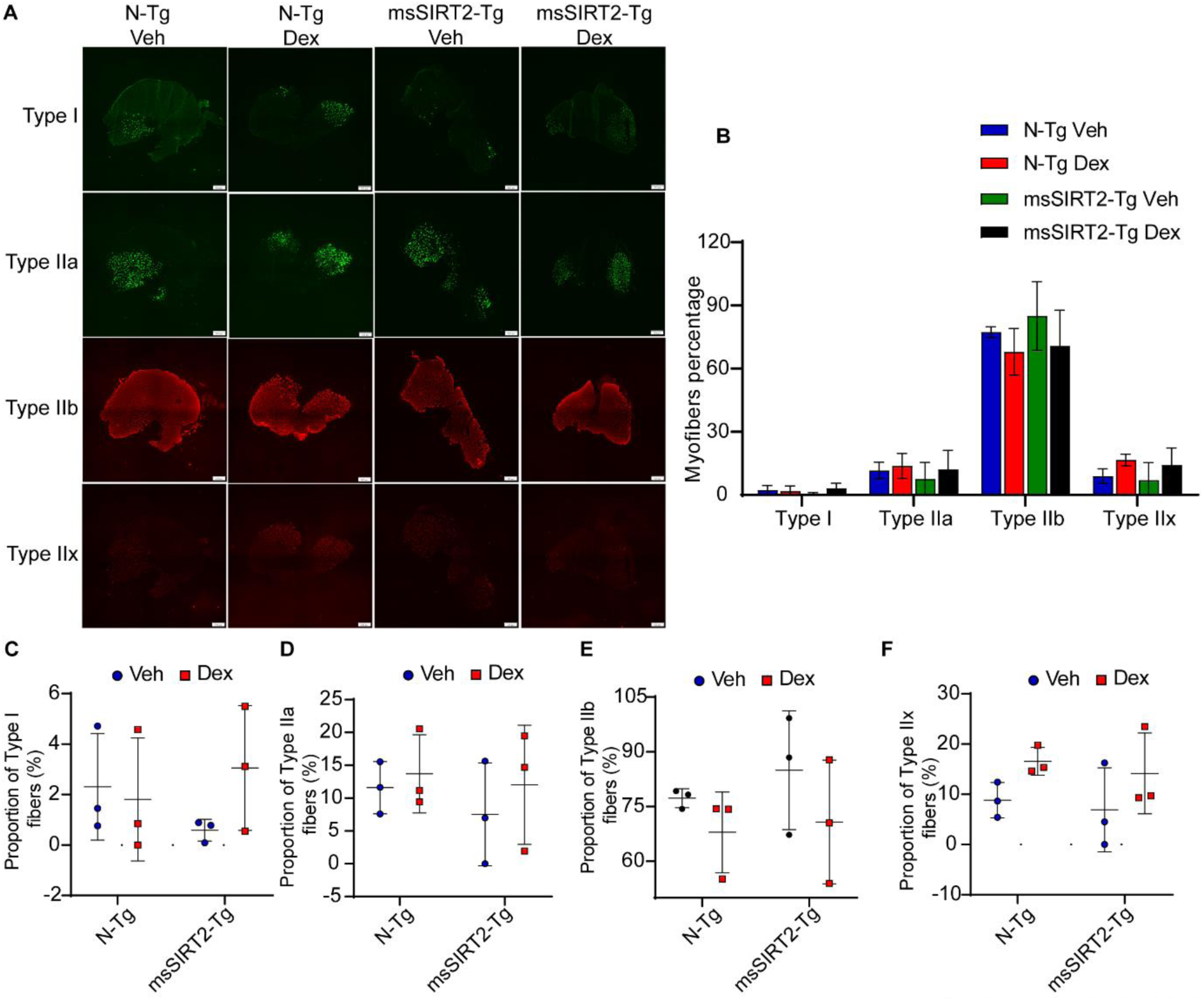
Muscle fibre type distribution in msSIRT2 transgenic mice **(A)** Representative immunofluorescence images showing various muscle fiber types (Type I, IIa (green) and Type IIb, IIx (red). Antibodies for Type I, IIa, IIb, IIx were probed in the gastrocnemius sections of non-transgenic control and msSIRT2 transgenic mice administered with either vehicle or dex(10mg/kg/day). Scale bar = 200 µm. **(B)** Bar graph showing the percentage of various fiber types in the gastrocnemius muscles of non-transgenic control and msSIRT2 transgenic mice injected with dex. Data represented as mean ± s.d., n = 3-4. **(C)-(F)** Scatterplot graphs representing the percentage of type I, IIa, IIb, and IIx as shown in Figure S7A in the gastrocnemius muscles of non-transgenic control and msSIRT2 transgenic mice administered with dex. Data represented as mean ± s.d., n = 3-4. Two-way ANOVA with Tukey’s multiple comparisons was used for statistical analysis (C, D, E, F).

**Supplementary figure 8.**
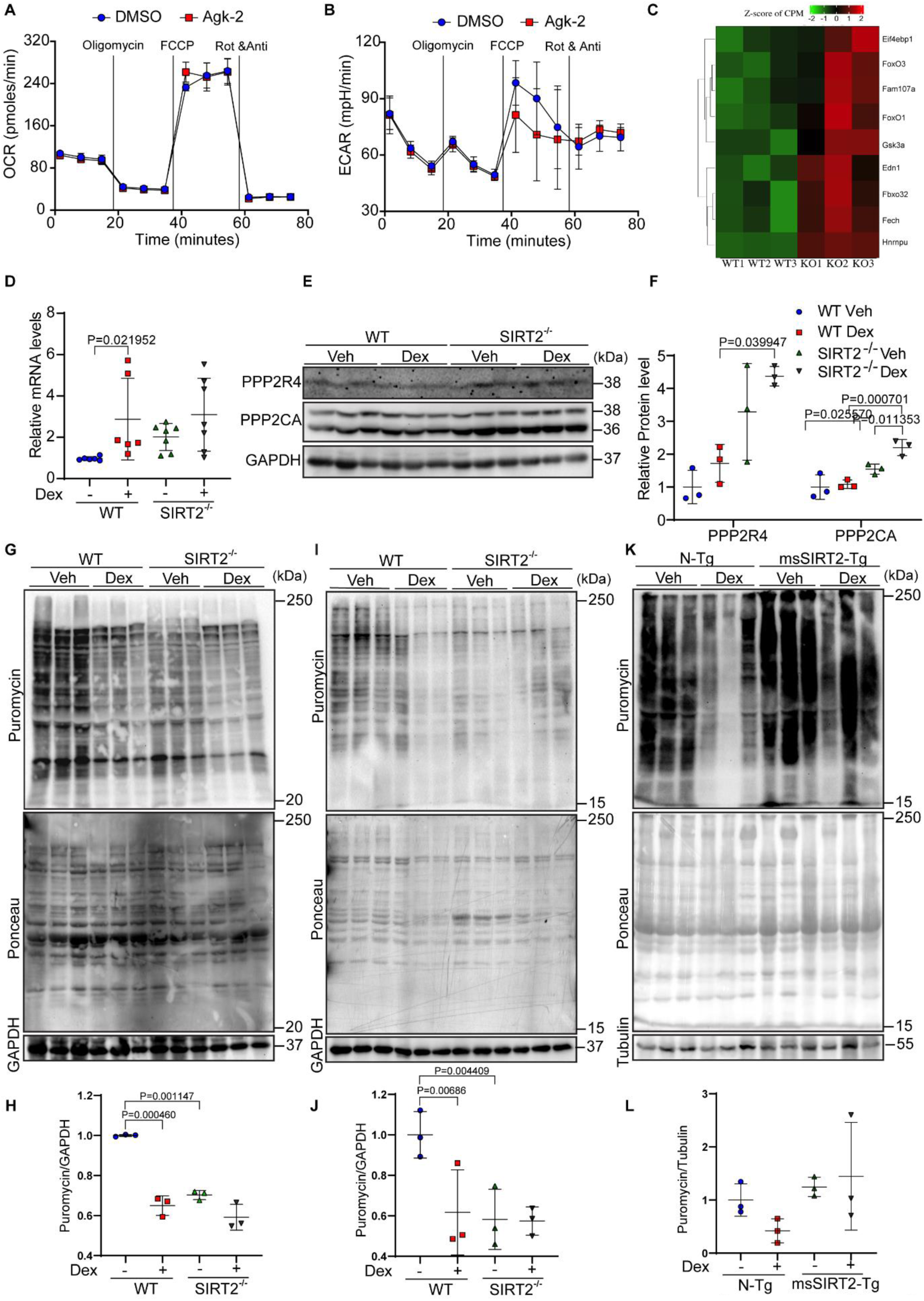
Characterization of oxidative metabolism and protein synthesis in SIRT2^-/-^ mice **(A)** Mitostress test representing OXPHOS-associated Oxygen Consumption Rate (OCR) in the DMSO and AGK2-treated primary myotubes. Data represented as mean ± s.d., *N*=3, *n* = 3. **(B)** Extracellular Acidification Rate (ECAR) associated with glycolysis in the DMSO and AGK2-treated primary myotubes. Data represented as mean ± s.d., *N*=3, *n* = 3. **(C)** Heatmap depicting significantly upregulated genes in SIRT2^-/-^ gastrocnemius muscles that are related to skeletal muscle atrophy. Upregulated and downregulated genes are represented in red and green, respectively. **(D)** qRT-PCR analysis for the relative expression of PP2A in dex-treated SIRT2^-/-^ mice, values are represented as fold change when normalized with vehicle treated wild-type control mice. Data represented as mean ± s.d., *n* = 6-7. **(E)** Representative immunoblot to check PPP2R4 and PPP2CA levels in SIRT2^-/-^ mice treated with Dex. Indicated sizes are in kilodaltons (kDa). **(F)** Quantification of PPP2R4 and PPP2CA levels in the gastrocnemius muscles of dex-treated SIRT2^-/-^ mice of Figure S 8E. Data represented as mean ± s.d., *n* = 3. **(G)** Representative immunoblot analysis depicting the changes in protein synthesis rates (tracked using SUnSET assay) in dex-treated SIRT2^-/-^ mice (male). **(H)** Quantitative representation of puromycin incorporation depicted in Figure S8(G). The results are expressed as the fold change relative to wild-type controls. Data represented as mean ± s.d., *n* = 3. **(I)** Representative immunoblot analysis depicting the changes in protein synthesis rates (tracked using SUnSET assay) in dex-treated SIRT2^-/-^ mice (female). **(J)** Quantitative representation of puromycin incorporation depicted in Figure S 8I. The results are expressed as the fold change relative to wild-type controls. Data represented as mean ± s.d., *n* = 3. **(K)** Representative immunoblot analysis depicting the changes in protein synthesis rates (tracked using SUnSET assay) in dex-treated muscle specific SIRT2 transgenic mice (male). **(L)** Quantitative representation of puromycin incorporation depicted in Figure S 8K. The results are expressed as the fold change relative to non-transgenic controls. Data represented as mean ± s.d., *n* = 3. Kruskal-Wallis test with Dunn’s multiple comparisons (D) Two-way ANOVA with Tukey’s multiple comparisons was used for statistical analysis (F, H, J, L).

**Supplementary figure 9.**
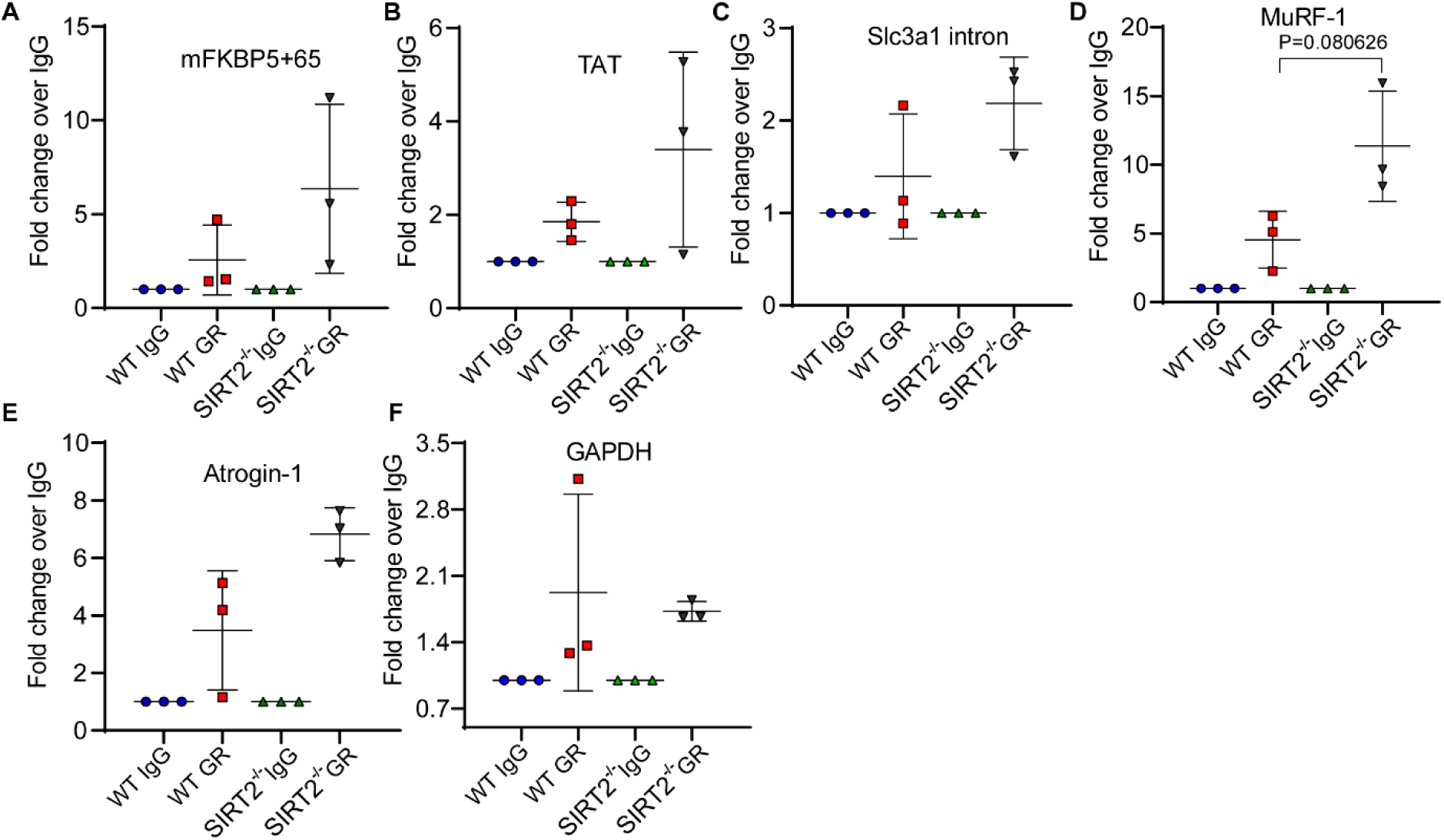
GR occupancy at GR target promoters (FKBP5, TAT, Slc3a1 intron) and FOXO3A target promoters (Atrogin-1 and MuRF-1) in SIRT2^-/-^ muscles **(A)-(F)** ChIP analysis showing GR binding on the promoters of indicated genes in gastrocnemius muscles of WT and SIRT2^-/-^ mice. Data are presented as mean ± s.d., n = 3. The two-tailed unpaired student’s t-test with Welch’s correction (between WT GR and KO GR groups) was used for statistical analysis (A, B, C, D, E, F)

**Supplementary figure 10.**
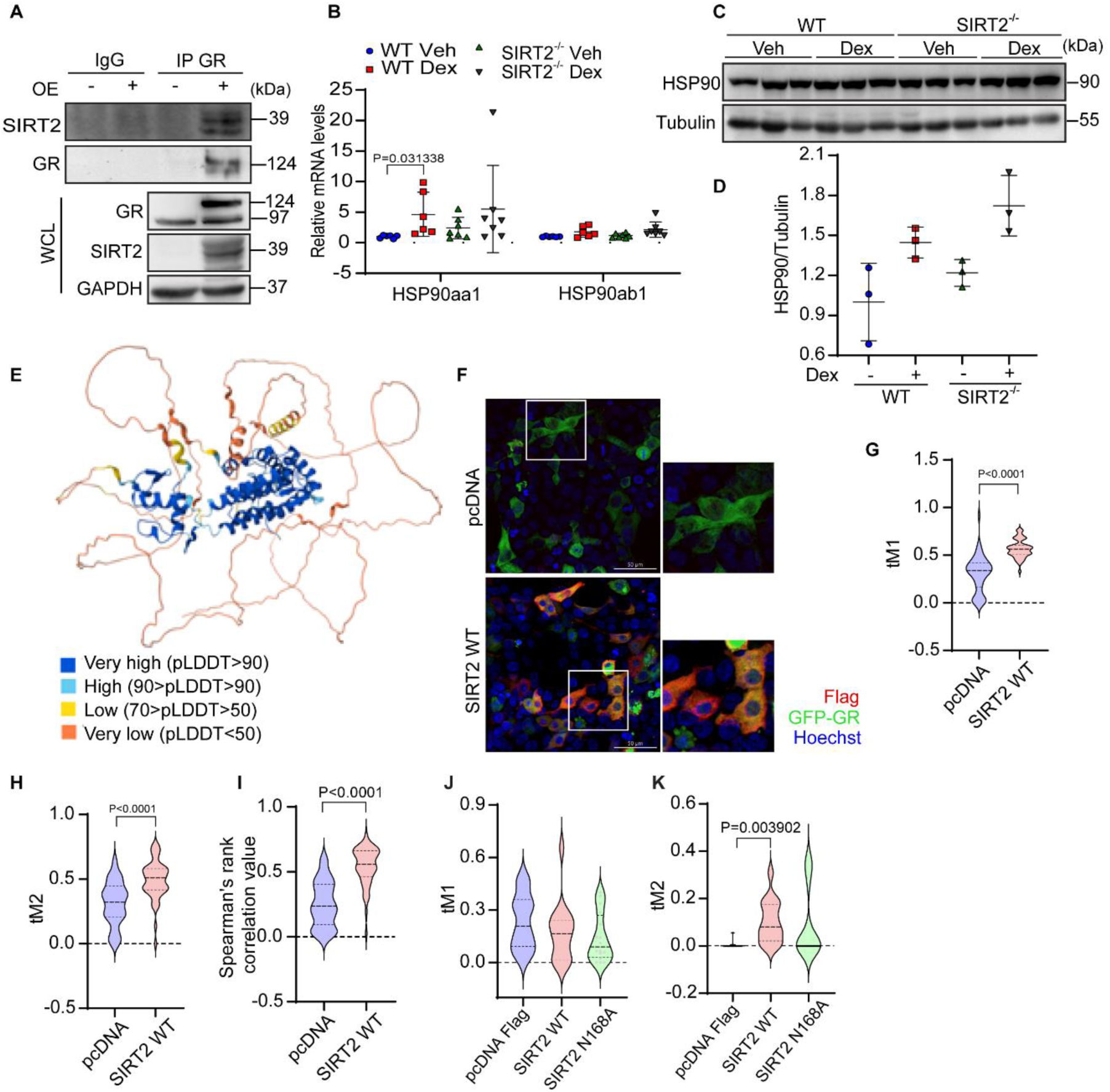
Characterisation of SIRT2-GR and SIRT2-HSP90 interaction *in vitro*. **(A)** Representative immunoblot images showing the interaction of SIRT2 with GR. GR was overexpressed in HEK293T cells, and GR was immunoprecipitated and probed for SIRT2. Whole cell lysate (WCL) was also immunoblotted to check the loading. Indicated band sizes are in kilodaltons (kDa). **(B)** qRT-PCR analysis for the relative expression of HSP90aa1 and HSP90ab1 in dex-treated SIRT2^-/-^ mice. Data represented as mean ± s.d., *n* = 6-7. **(C)** Representative immunoblot to check HSP90 levels in SIRT2^-/-^ mice treated with Dex. Indicated sizes are in kilodaltons (kDa). **(D)** Quantification of HSP90 levels in the gastrocnemius muscles of dex-treated SIRT2^-/-^ mice of Figure S 10C. Data represented as mean ± s.d., *n* = 3. **(E)** Representative predicted AlphaFold structure of GR in open conformation *(AF-P04150-F1-v4)*. **(F)** Representative confocal microscopy images showing the interaction of SIRT2 with GR upon overexpression of WT SIRT2 and GFP GR. **(G)** Quantification of figure S10(F) depicting threshold Manders coefficient1 (tM1). Fraction of signal from channel 1 (flag) that overlaps with signal from channel 2 (GFP-GR), after thresholding to exclude background. **(H)** Quantification of figure S10(F) depicting threshold Manders coefficient2 (tM2). Fraction of signal from channel 2 (GFP-GR) that overlaps with signal from channel 2 (flag), after thresholding to exclude background. **(I)** Quantification of figure S10(F) depicting Spearman’s rank correlation value. **(J)** Quantification of figure 7(F) depicting threshold Manders coefficient1 (tM1). Fraction of signal from channel 1 (flag) that overlaps with signal from channel 2 (GFP-GR), after thresholding to exclude background. **(K)** Quantification of figure 7(F) depicting threshold Manders coefficient2 (tM2). Fraction of signal from channel 2 (GFP-GR) that overlaps with signal from channel 2 (flag), after thresholding to exclude background. The two-tailed unpaired student’s t-test with Welch’s correction (H) Kruskal-Wallis test with Dunn’s multiple comparisons (B, J, K) Two-way ANOVA with Tukey’s multiple comparisons (D) Two-tailed Mann-Whitney statistical test (G, I) was used for statistical analysis.

## References

1. Frontera, W.R. and J. Ochala, Skeletal muscle: a brief review of structure and function. Calcif Tissue Int, 2015. 96(3): p. 183–95.

2. Allen, D.L., R.R. Roy, and V.R. Edgerton, Myonuclear domains in muscle adaptation and disease. Muscle Nerve, 1999. 22(10): p. 1350–60.

3. Ventadour, S. and D. Attaix, Mechanisms of skeletal muscle atrophy. Curr Opin Rheumatol, 2006. 18(6): p. 631–5.

4. Anker, S.D., et al., Wasting as independent risk factor for mortality in chronic heart failure. Lancet, 1997. 349(9058): p. 1050–3.

5. de Oliveira Nunes Teixeira, V., et al., Muscle wasting in collagen-induced arthritis and disuse atrophy. Exp Biol Med (Maywood), 2013. 238(12): p. 1421–30.

6. Wiedmer, P., et al., Sarcopenia - Molecular mechanisms and open questions. Ageing Res Rev, 2021. 65: p. 101200.

7. Hauser, C.A., M.R. Stockler, and M.H. Tattersall, Prognostic factors in patients with recently diagnosed incurable cancer: a systematic review. Support Care Cancer, 2006. 14(10): p. 999–1011.

8. Bjornholm, M. and J.R. Zierath, Insulin signal transduction in human skeletal muscle: identifying the defects in Type II diabetes. Biochem Soc Trans, 2005. 33(Pt 2): p. 354–7.

9. Diesel, W., et al., Morphologic features of the myopathy associated with chronic renal failure. Am J Kidney Dis, 1993. 22(5): p. 677–84.

10. Frimel, T.N., et al., A model of muscle atrophy using cast immobilization in mice. Muscle Nerve, 2005. 32(5): p. 672–4.

11. Vainshtein, A. and M. Sandri, Signaling Pathways That Control Muscle Mass. Int J Mol Sci, 2020. 21(13).

12. Damas, F., C.A. Libardi, and C. Ugrinowitsch, The development of skeletal muscle hypertrophy through resistance training: the role of muscle damage and muscle protein synthesis. Eur J Appl Physiol, 2018. 118(3): p. 485–500.

13. Zdychova, J. and R. Komers, Emerging role of Akt kinase/protein kinase B signaling in pathophysiology of diabetes and its complications. Physiol Res, 2005. 54(1): p. 1–16.

14. Whiteman, E.L., H. Cho, and M.J. Birnbaum, Role of Akt/protein kinase B in metabolism. Trends Endocrinol Metab, 2002. 13(10): p. 444–51.

15. White, J.P., et al., Testosterone regulation of Akt/mTORC1/FoxO3a signaling in skeletal muscle. Mol Cell Endocrinol, 2013. 365(2): p. 174–86.

16. Jensen, J., et al., Effects of adrenaline on whole-body glucose metabolism and insulin-mediated regulation of glycogen synthase and PKB phosphorylation in human skeletal muscle. Metabolism, 2011. 60(2): p. 215–26.

17. Beato, M., Gene regulation by steroid hormones. Cell, 1989. 56(3): p. 335–44.

18. Schakman, O., et al., Glucocorticoid-induced skeletal muscle atrophy. Int J Biochem Cell Biol, 2013. 45(10): p. 2163–72.

19. Kostyo, J.L. and A.F. Redmond, Role of protein synthesis in the inhibitory action of adrenal steroid hormones on amino acid transport by muscle. Endocrinology, 1966. 79(3): p. 531–40.

20. Shah, O.J., S.R. Kimball, and L.S. Jefferson, Acute attenuation of translation initiation and protein synthesis by glucocorticoids in skeletal muscle. Am J Physiol Endocrinol Metab, 2000. 278(1): p. E76–82.

21. Shah, O.J., S.R. Kimball, and L.S. Jefferson, Among translational effectors, p70S6k is uniquely sensitive to inhibition by glucocorticoids. Biochem J, 2000. 347(Pt 2): p. 389–97.

22. Liu, Z., et al., Glucocorticoids modulate amino acid-induced translation initiation in human skeletal muscle. Am J Physiol Endocrinol Metab, 2004. 287(2): p. E275–81.

23. Liu, Z., et al., Branched chain amino acids activate messenger ribonucleic acid translation regulatory proteins in human skeletal muscle, and glucocorticoids blunt this action. J Clin Endocrinol Metab, 2001. 86(5): p. 2136–43.

24. Hasselgren, P.O., Glucocorticoids and muscle catabolism. Curr Opin Clin Nutr Metab Care, 1999. 2(3): p. 201–5.

25. Bodine, S.C., et al., Identification of ubiquitin ligases required for skeletal muscle atrophy. Science, 2001. 294(5547): p. 1704–8.

26. Mitch, W.E. and A.L. Goldberg, Mechanisms of muscle wasting. The role of the ubiquitin-proteasome pathway. N Engl J Med, 1996. 335(25): p. 1897–905.

27. Khan, D., et al., SIRT6 transcriptionally regulates fatty acid transport by suppressing PPARgamma. Cell Rep, 2021. 35(9): p. 109190.

28. Matsushima, S. and J. Sadoshima, The role of sirtuins in cardiac disease. American Journal of Physiology-Heart and Circulatory Physiology, 2015. 309(9): p. H1375–H1389.

29. Dai, H., et al., Sirtuin activators and inhibitors: Promises, achievements, and challenges. Pharmacol Ther, 2018. 188: p. 140–154.

30. Lamming, D.W., et al., HST2 mediates SIR2-independent life-span extension by calorie restriction. Science, 2005. 309(5742): p. 1861–1864.

31. Wang, F., et al., SIRT2 deacetylates FOXO3a in response to oxidative stress and caloric restriction. Aging cell, 2007. 6(4): p. 505–514.

32. Wang, F. and Q. Tong, SIRT2 suppresses adipocyte differentiation by deacetylating FOXO1 and enhancing FOXO1’s repressive interaction with PPARgamma. Mol Biol Cell, 2009. 20(3): p. 801–8.

33. Sarikhani, M., et al., SIRT2 deacetylase represses NFAT transcription factor to maintain cardiac homeostasis. J Biol Chem, 2018. 293(14): p. 5281–5294.

34. Han, Z., et al., Role of SIRT2 in regulating the dexamethasone-activated autophagy pathway in skeletal muscle atrophy. Biochem Cell Biol, 2021. 99(5): p. 562–569.

35. Arora, A. and C.S. Dey, SIRT2 negatively regulates insulin resistance in C2C12 skeletal muscle cells. Biochim Biophys Acta, 2014. 1842(9): p. 1372–8.

36. Kadmiel, M. and J.A. Cidlowski, Glucocorticoid receptor signaling in health and disease. Trends Pharmacol Sci, 2013. 34(9): p. 518–30.

37. Pereira, R.M.R. and J.F. de Carvalho, Glucocorticoid-induced myopathy. Joint Bone Spine, 2011. 78(1): p. 41–44.

38. Fappi, A., et al., Skeletal Muscle Response to Deflazacort, Dexamethasone and Methylprednisolone. Cells, 2019. 8(5).

39. Livingstone, I., M.A. Johnson, and F.L. Mastaglia, Effects of dexamethasone on fibre subtypes in rat muscle. Neuropathol Appl Neurobiol, 1981. 7(5): p. 381–98.

40. Fappi, A., et al., Skeletal Muscle Response to Deflazacort, Dexamethasone and Methylprednisolone. Cells, 2019. 8(5).

41. North, B.J. and E. Verdin, Interphase nucleo-cytoplasmic shuttling and localization of SIRT2 during mitosis. PLoS One, 2007. 2(8): p. e784.

42. Pereira, J.M., et al., Infection Reveals a Modification of SIRT2 Critical for Chromatin Association. Cell Rep, 2018. 23(4): p. 1124–1137.

43. Verma, A., et al., Glutaredoxin 1 Downregulation in the Substantia Nigra Leads to Dopaminergic Degeneration in Mice. Mov Disord, 2020. 35(10): p. 1843–1853.

44. Kuo, T., et al., Genome-wide analysis of glucocorticoid receptor-binding sites in myotubes identifies gene networks modulating insulin signaling. Proc Natl Acad Sci U S A, 2012. 109(28): p. 11160–5.

45. Kedlian, V.R., et al., Human skeletal muscle aging atlas. Nat Aging, 2024. 4(5): p. 727–744.

46. Wang, Z., et al., Modulation of glucocorticoid receptor phosphorylation and transcriptional activity by a C-terminal-associated protein phosphatase. Mol Endocrinol, 2007. 21(3): p. 625–34.

47. DeFranco, D.B., et al., Protein phosphatase types 1 and/or 2A regulate nucleocytoplasmic shuttling of glucocorticoid receptors. Mol Endocrinol, 1991. 5(9): p. 1215–28.

48. Jagoe, R.T., et al., Skeletal muscle mRNA levels for cathepsin B, but not components of the ubiquitin-proteasome pathway, are increased in patients with lung cancer referred for thoracotomy. Clin Sci (Lond), 2002. 102(3): p. 353–61.

49. Almon, R.R., et al., Microarray analysis of the temporal response of skeletal muscle to methylprednisolone: comparative analysis of two dosing regimens. Physiol Genomics, 2007. 30(3): p. 282–99.

50. Komamura, K., et al., Differential gene expression in the rat skeletal and heart muscle in glucocorticoid-induced myopathy: analysis by microarray. Cardiovasc Drugs Ther, 2003. 17(4): p. 303–10.

51. Lecker, S.H., et al., Multiple types of skeletal muscle atrophy involve a common program of changes in gene expression. FASEB J, 2004. 18(1): p. 39–51.

52. Waddell, D.S., et al., The glucocorticoid receptor and FOXO1 synergistically activate the skeletal muscle atrophy-associated MuRF1 gene. Am J Physiol Endocrinol Metab, 2008. 295(4): p. E785–97.

53. Dobson, M., et al., Bimodal regulation of FoxO3 by AKT and 14-3-3. Biochim Biophys Acta, 2011. 1813(8): p. 1453–64.

54. Brunet, A., et al., Akt promotes cell survival by phosphorylating and inhibiting a Forkhead transcription factor. Cell, 1999. 96(6): p. 857–68.

55. Kuo, T., et al., Transcriptional regulation of FoxO3 gene by glucocorticoids in murine myotubes. Am J Physiol Endocrinol Metab, 2016. 310(7): p. E572–85.

56. Lützner, N., et al., FOXO3 is a glucocorticoid receptor target and regulates LKB1 and its own expression based on cellular AMP levels via a positive autoregulatory loop. PLoS One, 2012. 7(7): p. e42166.

57. Bolster, D.R., et al., AMP-activated protein kinase suppresses protein synthesis in rat skeletal muscle through down-regulated mammalian target of rapamycin (mTOR) signaling. J Biol Chem, 2002. 277(27): p. 23977–80.

58. Williamson, D.L., et al., Time course changes in signaling pathways and protein synthesis in C2C12 myotubes following AMPK activation by AICAR. Am J Physiol Endocrinol Metab, 2006. 291(1): p. E80–9.

59. Jaitovich, A., et al., High CO2 levels cause skeletal muscle atrophy via AMP-activated kinase (AMPK), FoxO3a protein, and muscle-specific Ring finger protein 1 (MuRF1). J Biol Chem, 2015. 290(14): p. 9183–94.

60. Baehr, L.M., J.D. Furlow, and S.C. Bodine, Muscle sparing in muscle RING finger 1 null mice: response to synthetic glucocorticoids. J Physiol, 2011. 589(Pt 19): p. 4759–76.

61. Nader, N., G.P. Chrousos, and T. Kino, Circadian rhythm transcription factor CLOCK regulates the transcriptional activity of the glucocorticoid receptor by acetylating its hinge region lysine cluster: potential physiological implications. FASEB J, 2009. 23(5): p. 1572–83.

62. Sun, K., et al., SIRT2 suppresses expression of inflammatory factors via Hsp90-glucocorticoid receptor signalling. J Cell Mol Med, 2020. 24(13): p. 7439–7450.

63. Franco, P.J., et al., The orphan nuclear receptor TR2 interacts directly with both class I and class II histone deacetylases. Mol Endocrinol, 2001. 15(8): p. 1318–28.

64. Ishii, S., The Role of Histone Deacetylase 3 Complex in Nuclear Hormone Receptor Action. Int J Mol Sci, 2021. 22(17).

65. Weikum, E.R., X. Liu, and E.A. Ortlund, The nuclear receptor superfamily: A structural perspective. Protein Sci, 2018. 27(11): p. 1876–1892.

66. Wu, W., et al., Glucocorticoid receptor activation signals through forkhead transcription factor 3a in breast cancer cells. Mol Endocrinol, 2006. 20(10): p. 2304–14.

67. Wang, F., et al., SIRT2 deacetylates FOXO3a in response to oxidative stress and caloric restriction. Aging Cell, 2007. 6(4): p. 505–14.

68. Hu, Z., et al., Endogenous glucocorticoids and impaired insulin signaling are both required to stimulate muscle wasting under pathophysiological conditions in mice. J Clin Invest, 2009. 119(10): p. 3059–69.

69. Shimizu, N., et al., Crosstalk between Glucocorticoid Receptor and Nutritional Sensor mTOR in Skeletal Muscle. Cell Metabolism, 2011. 13(2): p. 170–182.

70. Wang, Z., J. Frederick, and M.J. Garabedian, Deciphering the phosphorylation “code” of the glucocorticoid receptor in vivo. J Biol Chem, 2002. 277(29): p. 26573–80.

71. Itoh, M., et al., Nuclear export of glucocorticoid receptor is enhanced by c-Jun N-terminal kinase-mediated phosphorylation. Mol Endocrinol, 2002. 16(10): p. 2382–92.

72. Bruna, A., et al., Glucocorticoid receptor-JNK interaction mediates inhibition of the JNK pathway by glucocorticoids. Embo j, 2003. 22(22): p. 6035–44.

73. Sarikhani, M., et al., SIRT2 regulates oxidative stress-induced cell death through deacetylation of c-Jun NH(2)-terminal kinase. Cell Death Differ, 2018. 25(9): p. 1638–1656.

74. Ravi, V., et al., Measuring Protein Synthesis in Cultured Cells and Mouse Tissues Using the Non-radioactive SUnSET Assay. Curr Protoc Mol Biol, 2020. 133(1): p. e127.

75. Tamta, A.K., et al., Cultured Neonatal Murine Primary Myotubes as a Model to Study Muscle Atrophy. Curr Protoc, 2022. 2(11): p. e616.

76. Ross, J.M., Visualization of mitochondrial respiratory function using cytochrome c oxidase/succinate dehydrogenase (COX/SDH) double-labeling histochemistry. J Vis Exp, 2011(57): p. e3266.

77. Sato, S., et al., TWEAK promotes exercise intolerance by decreasing skeletal muscle oxidative phosphorylation capacity. Skelet Muscle, 2013. 3(1): p. 18.

78. Wickham, H., et al., Welcome to the Tidyverse. Journal of Open Source Software, 2019. 4: p. 1686.

79. Zhou, Y., et al., Metascape provides a biologist-oriented resource for the analysis of systems-level datasets. Nat Commun, 2019. 10(1): p. 1523.

80. Gu, Z., R. Eils, and M. Schlesner, Complex heatmaps reveal patterns and correlations in multidimensional genomic data. Bioinformatics, 2016. 32(18): p. 2847–9.

81. Stevenson, E.J., et al., Global analysis of gene expression patterns during disuse atrophy in rat skeletal muscle. J Physiol, 2003. 551(Pt 1): p. 33–48.

82. Kedia, S., et al., Real-time nanoscale organization of amyloid precursor protein. Nanoscale, 2020. 12(15): p. 8200–8215.

83. Rajeev, P., et al., Nanoscale regulation of Ca(2+) dependent phase transitions and real-time dynamics of SAP97/hDLG. Nat Commun, 2022. 13(1): p. 4236.

84. Nair, D., et al., Super-resolution imaging reveals that AMPA receptors inside synapses are dynamically organized in nanodomains regulated by PSD95. J Neurosci, 2013. 33(32): p. 13204–24.

85. Spurthi, K.M., et al., Toll-like receptor 2 deficiency hyperactivates the FoxO1 transcription factor and induces aging-associated cardiac dysfunction in mice. J Biol Chem, 2018. 293(34): p. 13073–13089.

86. Hudson, W.H., et al., Cryptic glucocorticoid receptor-binding sites pervade genomic NF-κB response elements. 2018. 9(1): p. 1–13.

87. Warnecke, A., et al., PyTMs: a useful PyMOL plugin for modeling common post-translational modifications. 2014. 15(1): p. 1–12.

88. Spoel, V.D.J.Z., GROMACS 2020.2 Source code.

89. Pettersen, E.F., et al., UCSF Chimera—a visualization system for exploratory research and analysis. 2004. 25(13): p. 1605–1612.

90. Grant, B.J., L. Skjærven, and X.Q.J.P.S. Yao, The Bio3D packages for structural bioinformatics. 2020.

